# Gene flux and acid-imposed selection are the main drivers of antimicrobial resistance in broiler chicks infected with *Salmonella enterica* serovar Heidelberg

**DOI:** 10.1101/2021.02.25.432983

**Authors:** Adelumola Oladeinde, Zaid Abdo, Maximilian O. Press, Kimberly Cook, Nelson A. Cox, Benjamin Zwirzitz, Reed Woyda, Steven M. Lakin, Jesse C. Thomas, Torey Looft, Douglas E. Cosby, Arthur Hinton, Jean Guard, Eric Line, Michael J. Rothrock, Mark E. Berrang, Kyler Herrington, Gregory Zock, Jodie Plumblee Lawrence, Denice Cudnik, Sandra House, Kimberly Ingram, Leah Lariscy, Martin Wagner, Samuel E. Aggrey, Lilong Chai, Casey Ritz

## Abstract

Antimicrobial resistance (AR) spread is a worldwide health challenge, stemming in large part, from the ability of microbes to share their genetic material through horizontal gene transfer (HGT). Overuse and misuse of antibiotics in clinical settings and in food production have been linked to this increased prevalence and spread of AR. Consequently, public health and consumer concerns have resulted in a remarkable recent reduction in antibiotics used for food animal production. This is driven by the assumption that removing this selective pressure will favor the recovery of antibiotic susceptible taxa and will limit AR sharing through HGT, allowing the currently available antibiotic arsenal to be effective for a longer period. In this study we used broiler chicks raised antibiotic-free and *Salmonella enterica* serovar Heidelberg (SH), as a model food pathogen, to test this hypothesis. Our results show that neonatal broiler chicks challenged with an antibiotic susceptible SH strain and raised without antibiotics carried susceptible and multidrug resistance SH strains 14 days after challenge. SH infection perturbed the microbiota of broiler chicks and gavaged chicks acquired antibiotic resistant SH at a higher rate. We determined that the acquisition of a plasmid from commensal *Escherichia coli* population conferred multidrug resistance phenotype to SH recipients and carriage of this plasmid increased the fitness of SH under acidic selection pressure. These results suggest that HGT of AR shaped the evolution of SH and that antibiotic use reduction alone is insufficient to limit antibiotic resistance transfer from commensal bacteria to *Salmonella*.

**Importance:** The reported increase in antibiotic resistant bacteria in humans have resulted in a major shift away from antibiotics use in food animal production. This has been driven by the assumption that removing antibiotics will select for antibiotic susceptible bacterial taxa, and this in turn will allow the currently available antibiotic arsenal to be more effective. This shift in practice has highlighted new questions that need to be answered to assess the effectiveness of antibiotic removal in reducing the spread of antibiotic resistance bacteria. This research demonstrates that antibiotic susceptible *Salmonella* Heidelberg strains can acquire multidrug resistance from commensal bacteria present in the gut of neonatal broiler chicks, even in the absence of antibiotic selection. We demonstrate that exposure to acidic pH drove the horizontal transfer of antimicrobial resistance plasmids and suggests that simply removing antibiotics from food-animal production might not be sufficient to limit the spread of antimicrobial resistance.

## Introduction

Antimicrobial resistance (AR) spread is a worldwide health challenge^1, 2^. A major aspect of this challenge stems from the ability of microbes to share their genetic material through horizontal gene transfer (HGT)^3^. Exposure to antibiotics can select for a microbiota of commensals and pathogens that can withstand antibiotic selective pressure, creating a potential reservoir of resistance genes that are shared with susceptible taxa allowing for their survival and proliferation^3^. Overuse and misuse of antibiotics in a clinical setting^4^ and in food production^5^ have been shown to result in such selective environments contributing to the increased prevalence and spread of AR. These public health concerns have resulted in a major shift away from antibiotics use in food production. This was driven by the assumption that removing this selective pressure will favor the recovery of susceptible taxa, and that this in turn will allow the currently available antibiotic arsenal to be effective for a longer period^6–9^. This shift in food production practices has highlighted new questions that need to be answered to understand and assess the effectiveness of antibiotic removal in reducing the spread and prevalence of AR in food borne pathogens. There has been a remarkable reduction in antibiotics sold or distributed in the United States (US) for use in food-producing animals since the publication of the US Food and Drug Administration’s guidance for industry #209 (2012) and #213 (2013) and the Veterinary Feed Directive^10^. Chicken production accounted for the largest reduction in medically and nonmedically important antibiotics sold (∼47% from 2016 to 2017), also driven by consumer concerns over food safety related to antibiotics use^11^. *Salmonella enterica* serovar Heidelberg (SH) strain is one of the prolific serovars causing salmonellosis in the US and poultry is by far the major vector for SH infections^12–14^. One of the largest food-borne outbreaks in US history was associated with the consumption of chicken contaminated with SH^15^. During this outbreak, SH was present in blood samples of case-patients (15%) and resulted in an overall hospitalization rate of 38% (200 case-patients). Poultry and clinical strains of SH can be resistant to at least 1 antibiotic in 3 or more drug classes. Therefore, we focused on studying antibiotic-free broiler chicken production, the most consumed meat in the US, with a goal to evaluate the prevalence of AR and its potential transfer to SH, an important model pathogen.

Our results suggested that *Salmonella* can acquire multidrug resistance from the bacteria of neonatal broiler chicks raised without antibiotics. We determined that *E. coli* strains carrying IncI1 plasmids were the main reservoir of transferable AR in broiler chicks and confirmed that *E. coli* was the donor of IncI1-ST26 (pST26) carrying antibiotic, metal, and disinfectant genes to SH. The presence of pST26 did not impose a fitness cost to the SH host and exposure to acidic pH posed a selection pressure on pST26 plasmid population. Commensals like *E. coli* are a reservoir of mobile genetic elements including plasmids and bacteriophages. Therefore, we conclude that a simple reduction in antibiotic use might not be sufficient to prevent transfer of these mobile elements to a major food-borne pathogen like *Salmonella*.

## Results

### S. Heidelberg abundance in ceca and litter is influenced by the route of challenge

To investigate if the route of exposure to SH affected intestinal and litter colonization, we challenged 150 one day old broiler chicks with a nalidixic (nal) resistant strain of SH (SH-2813_nal_) either by oral gavage (OG), intracloacally (IC) or by the seeder (SB) method (i.e., a few chicks (n =5) were orally challenged and co-mingled with unchallenged chicks (n = 20)) (Fig. 1a). Chicks were grown for 14 days on fresh pine shavings in four separate floor pens (1.8 L ×1.16 W m or 5.9 L×3.8 W ft) including an unchallenged control (CTRL) group (n=25). Chicks were not administered any medication or antibiotics for the duration of the study. The challenge experiments, trial 1 and trial 2, were performed in September 2017 and April 2018, respectively and we have published the rate SH colonized the ceca^16^. SH concentration in the ceca of gavaged chicks (SB and OG) was higher than IC for trial 1 (H = 0.94, df =2, *P* = 0.62) and OG was higher than IC for trial 2 (H = 10.24, df =2, *P* = 0.006) (Fig. 1b). In the litter, SH levels were higher for IC pens compared to SB and OG in trial 1 and higher in OG pens compared to IC for trial 2 (Fig. 1c). The percentage of chicks that had SH detected in their ceca was 80% for OG and SB and 100% for IC during trial 1. For trial 2, 100% of the OG and IC were positive and 40% for SB^15^. The ceca and litter of CTRL were negative for SH by culture; however, one litter sample was positive after 24 h enrichment in buffered peptone water. Litter pH and moisture was higher for trial 1 (pH = 6.86 ± 0.27; moisture = 25± 0.03 %) compared to trial 2 (6.54 ± 0.17; moisture = 21± 0.03 %) (Extended data Fig. 1a and b).

**Figure 1.**
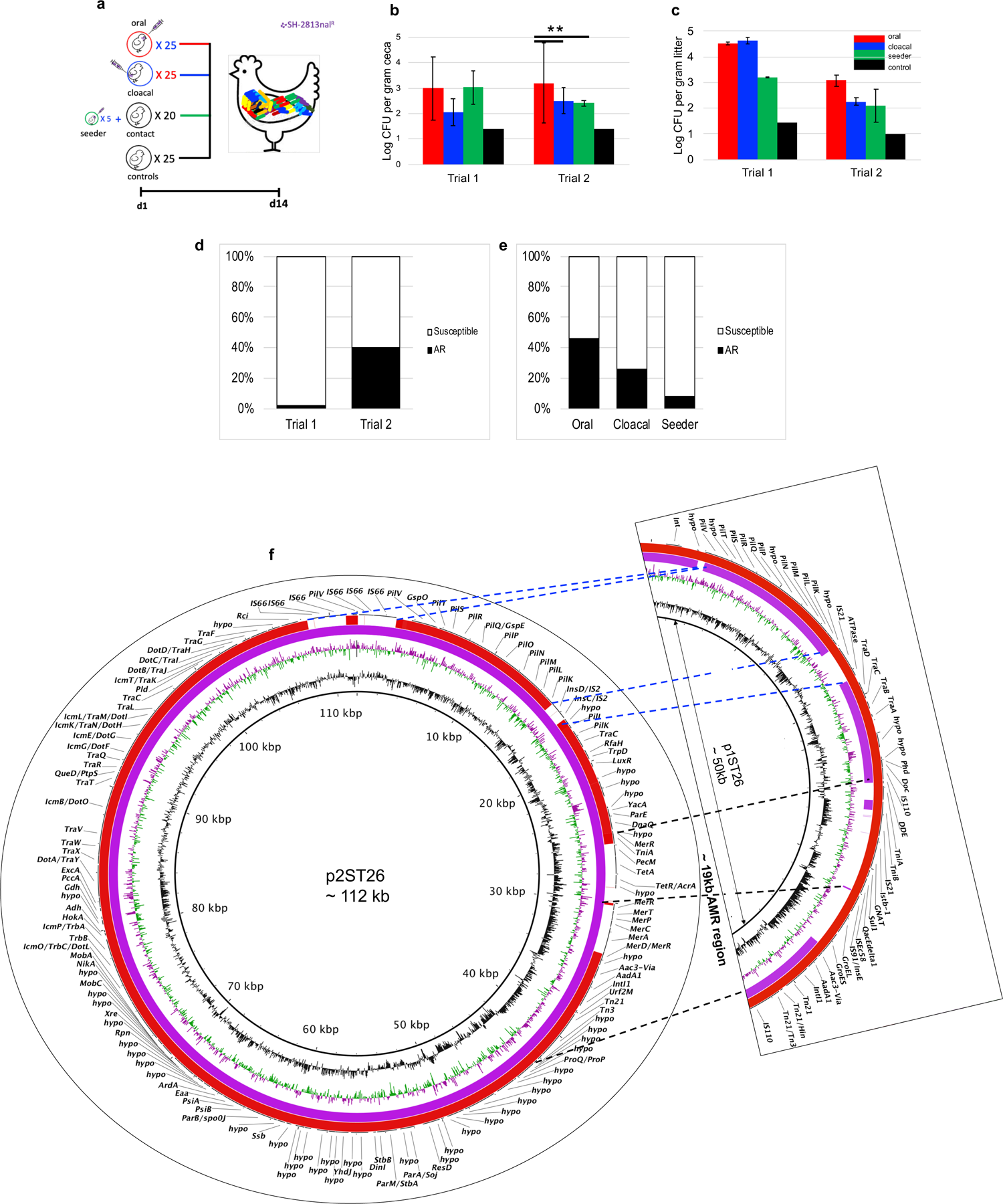
Broiler chicks challenged with an antibiotic susceptible strain of SH carried antibiotic resistance SH strains 2-weeks after challenge. **a**, Experimental design for trial 1 and trial 2 conducted in September 2017 and April 2018, respectively. **b,c** SH load in the ceca (n = 10 per trial) and litter (n = 2 per trial) 2 weeks after challenging 1-day broiler chicks (100 chicks/trial) with 10^6^ colony forming units (CFU) of SH by oral gavage, cloacal or through the seeder method (*P < 0.01 (Kruskal-Wallis test, multiple comparison correction; error bar = Standard deviation; limit of quantification = Log 1.3 CFU/g litter). **d**, Percentage of SH isolates that acquired antibiotic resistance in trial 1 (n = 92) and trial 2 (n = 158) **e**, Percentage of SH isolates in trial 2 that acquired antibiotic resistance grouped by the route the broiler chicks got challenged (n = 54 62, and 42 for oral, cloaca and seeder, respectively. **f,** BLASTn alignment of the IncI1-complex plasmid acquired during trial 1 (p1ST26) and trial 2 (p2ST26) – dashed black lines highlights mobile region encoding antimicrobial resistance genes while blue dashed lines shows other regions that are different between p1ST26 and p2ST26.

### S. Heidelberg perturbed the ceca and litter bacteria in a challenge route-specific manner

Next, we questioned if SH gut colonization perturbed changes in the ceca and litter microbiota. We profiled the microbial community present in the ceca and litter using 16S ribosomal RNA gene amplicon sequencing. The bacterial alpha diversity was higher for litter samples compared to the ceca for OG for all alpha diversity measures (Observed, Chao1, Shannon and InvSimpson) examined (*P* < 0.01) (Extended data Fig. 2a). In contrast, ceca from OG had lowest alpha diversity for all measures used compared to CTRL (*P* < 0.05). Observed and Chao1 alpha diversity indices were higher for litter compared to ceca for CTRL and IC (*P* = 0.007). Beta diversity differed between litter and ceca and the route of SH challenge affected the beta diversity of the litter (F = 43.7, df = 11, *P* = 0.001) more than the ceca (F = 2.21, df = 29 *P* = 0.007). Cecal samples were more dispersed than litter samples and OG chicks had the highest beta diversity (Fig. 2a). *Firmicutes*, *Proteobacteria*, *Actinobacteria* and *Bacteriodetes* were the dominant phyla present in litter (Fig. 2b). Litter from OG and SB carried higher abundance of *Bacteriodetes*, whereas IC and CTRL carried higher abundance of *Firmicutes* than OG and SB. *Firmicutes* dominated the cecal bacteria of broiler chicks and *Ruminococcaceae* and *Lachnospiraceae* (class *Clostridia*) were the major families. *Proteobacteria* was higher in the ceca of CTRL than OG or IC (X²= 0.6412, *P*=0.72) (Fig.2b). Amplicon sequence variants (ASV) matching *Subdolinagrum*, *Lactobacillus* and *Family_Lachnospiraceae* were the most abundant ASVs in OG and IC compared to CTRL in the ceca (Extended data Fig.2b). In contrast, *E. coli/Shigella* (*E. coli* and *Shigella* are not distinguishable by 16S rRNA gene sequencing) and *Klebsiella* were the most abundant ASVs in litter. *E. coli* was higher in IC and CTRL compared to OG and SB and *Klebsiella* was higher in CTRL compared to OG, IC and SB (*E. coli*: X²= 9.9744, *P*=0.018; *Klebsiella*: X²=9.6667, *P*=0.021). This result suggested that SH perturbed the bacterial community present in the litter and ceca of broiler chicks especially the *Enterobacteriaceae*.

**Figure 2.**
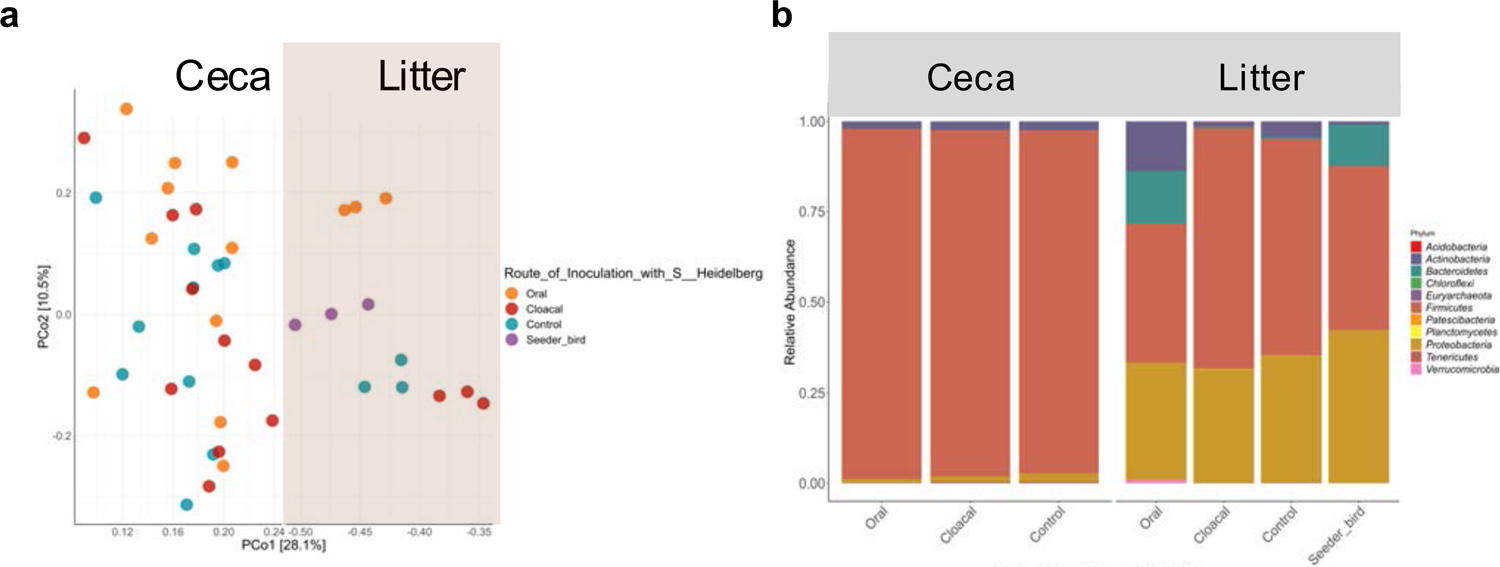
SH infection perturbed the bacterial community present in ceca and litter of broiler chickens. **a**, Principal coordinates analysis of Bray-Curtis distances based on 16S rRNA gene libraries obtained from ceca and litter samples. Each point represents values from individual libraries with colors expressing the route of inoculation of SH in for respective samples. b, Phylum-level classification of 16S rRNA gene sequence reads in ceca and litter grouped by route of SH infection. Data represent average of amplicon sequence variant (ASV) counts from replicate libraries for each category.

### S. Heidelberg developed multidrug resistance after IncI1 plasmid acquisition

We investigated if SH isolates developed AR after 14 days of challenge. We used the national antimicrobial resistance monitoring system sensititre ™ panel for Gram-negative bacteria to conduct antibiotic susceptibility testing (AST) on 250 SH isolates recovered from either the ceca or litter of challenged chicks. SH-2813_nal_ carries a *gyrA* mutation for nal resistance, therefore all isolates recovered were resistant to nal. SH also carries a chromosome-encoded *fosA7* gene that confers resistance to fosfomycin^17^. In trial 1, only 2 % of SH isolates (n = 92) developed resistance to an antibiotic. These two isolates were resistant to gentamicin, streptomycin and sulfisoxazole. For trial 2, 40 % of the isolates developed resistance to at least one antibiotic (Fig. 1d) and > 86 % of them were resistant to 2 or more drug classes like the isolates from trial 1 (Extended data Fig. 3a). However, trial 2 isolates developed AR to tetracyclines instead of sulfisoxazole. A disc diffusion assay on a selected number of multidrug resistant (MDR) isolates (n = 4) revealed that MDR isolates were also resistance to tobramycin, netilmicin and kanamycin (data not shown). Additionally, OG chicks carried a higher percentage (46 %) of MDR isolates than IC (24 %) or SB (8%) chicks (X^2^ = 29.2, df = 2, *P* < 0.001) (Fig. 1e).

**Figure 3.**
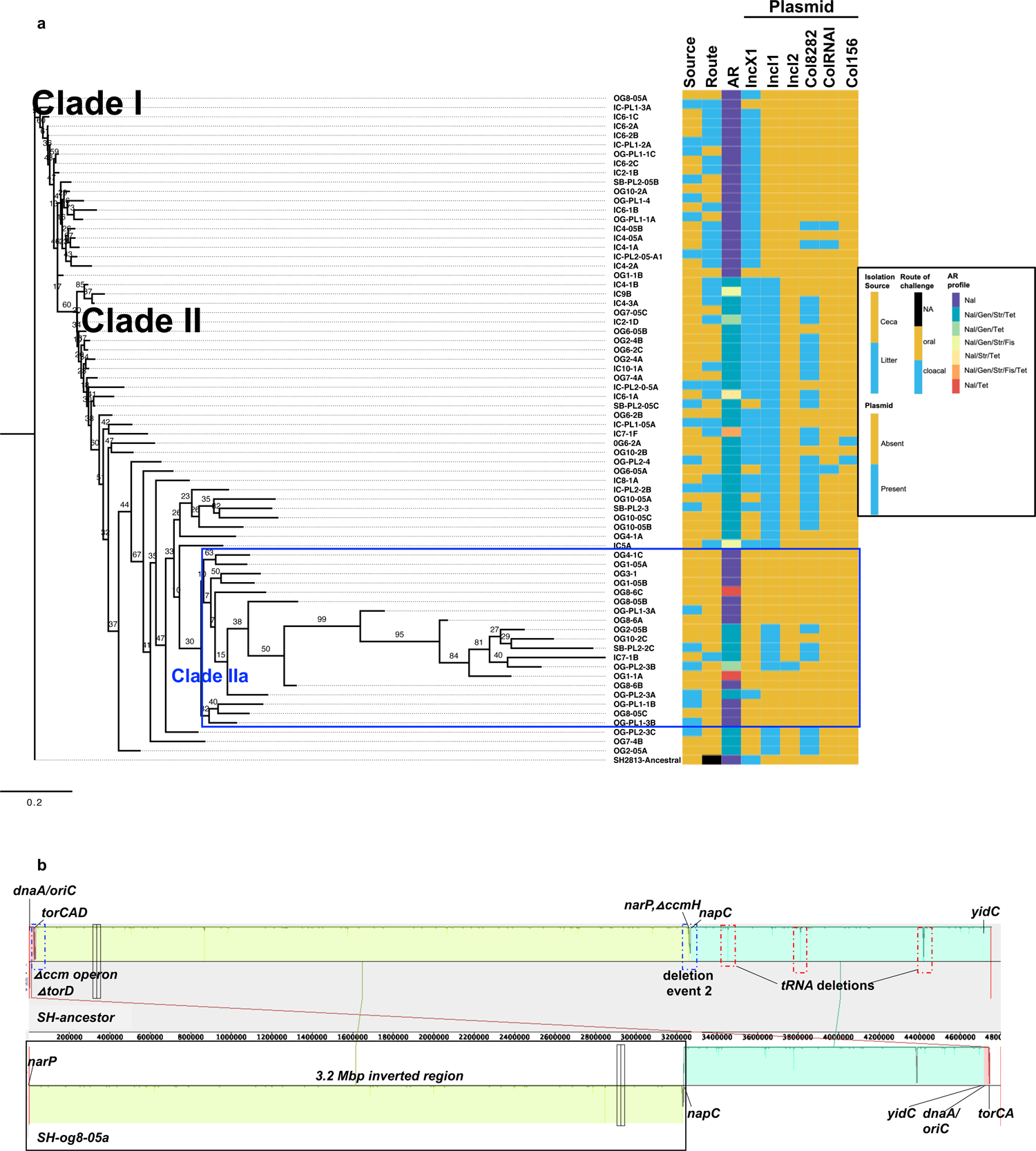
Horizontal gene transfer and genome inversion shaped the fitness of SH in broiler chicks. **a**, Maximum likelihood tree constructed using accessory genes present in SH isolates (n=72) recovered from the ceca and litter of broiler chicks colonized with SH. The GTR model of nucleotide substitution and the GAMMA model of rate heterogeneity were used for sequence evolution prediction. Numbers shown next to the branches represent the percentage of replicate trees where associated isolates cluster together based on ∼100 bootstrap replicates. Tree was rooted with the ancestral susceptible SH strain. Clade numbers were arbitrary assigned to facilitate discussion on isolates that acquired antibiotic resistance. Blue rectangular box highlights the clade with susceptible strains nested within antibiotic resistance strains due to sequencing bias and misassembly. (Legend: Nal – nalidixic acid, Gen – gentamicin, Str – streptomycin, Tet – tetracycline, Fis – sulfisoxazole, NA – Not applicable) **b,** Mauve visualization of the inverted genome of SH strain og8-05a. The chromosomal contigs of og8-05a was aligned and ordered to the complete chromosome of SH ancestor. A colored similarity plot is shown for each genome, the height of which is proportional to the level of sequence identity in that region. When the similarity plot points downward it indicates an alignment to the reverse strand of the SH ancestor genome i.e., inversion. Segment highlighted with solid black rectangular box shows the inverted region in og8-05a, while dashed blue and red rectangular boxes denotes segments with relevant mutations.

To find AR determinants responsible for the acquired AR phenotype, we performed whole genome sequencing (WGS) using illumina short read sequencing technology on sixty-nine SH isolates from trial 2 and two isolates from trial 1. We focused our resources on trial 2, for which AR acquisition was higher than trial 1. The isolates sequenced were either susceptible (n = 33) or antibiotic resistant (n = 38). The ancestral SH-2813_nal_^R^ (isolate used for challenge) harbors an IncX1 plasmid and carriage of this plasmid does not confer AR in SH. The IncX1 *inc* region was not detected by plasmidFinder^18^ in 42 of the 71 isolates sequenced. Thirty-nine percent of SH isolates acquired Col plasmids, but none carried a known antibiotic resistance gene (ARG). ARGs known to confer resistance to aminoglycosides (*aadA1, aac(3)-Via*) and tetracyclines (*tetA*) were detected in all AR isolates. The AR isolates from trial 1 harbored a sulphonamide resistance gene (*sul1*) in addition to aminoglycoside genes but no *tet* gene. The ARGs were found on a plasmid belonging to *inc* group IncI1 (Fig. 1f) and plasmid Multi Locus Sequence Type 26 (pST26). A change in AR profile was observed in four SB isolates - for these isolates, AST confirmed them to be MDR but after WGS, the AR determinants were not detected. Upon AST re-test, these isolates were found to be susceptible. Acquired ARG was absent in four isolates with resistance to either tetracycline or streptomycin.

### HGT and chromosome rearrangement shaped the genome of *S*. Heidelberg

After introduction of SH to the broiler gut microbiota, we hypothesized that successful colonization will be dependent on gene flux (gain and loss of genes) and mutations. To this end, we reconstructed a maximum-likelihood (ML) tree based on the pan genome and mutations of SH strains recovered after 2 weeks in the gut or litter. The core genome (genes present in ≥ 95% of the strains) and accessory genome (genes present in < 95% of the strains) composed of 3729 and 1531 genes, respectively. A total of 91 new mutations (SNPs and Indels) were on the chromosome of evolved strains and no single mutation was present in every strain. Rather, mutations were unique to individual strains or shared between 2-26 isolates (Supplementary data 1). The accessory genome tree grouped the strains based on AR phenotype, plasmid and ARG carried (Fig. 3a).

Contrastingly, the core genome and SNP-based trees did not provide a clear division of strains based on these characteristics and clades had low bootstrap support values (Extended data Fig. 3b and c). Acquisition of IncI (IncI1 and IncI2) plasmid, Col plasmids and the presence/absence of IncX1 plasmids defined the clades seen on the accessory genome tree. Susceptible strains carrying only IncX1 plasmids represented Clade I and MDR strains dominated Clade II. Clade II strains carried either IncI1 only or IncI1 plus a Col plasmid or both were present with an IncX1 plasmid (Fig. 3a).

A few susceptible and strains with no plasmid (n = 12) were nested within clade II and the genome of these strains exhibited signs of substantial gene loss (Fig. 3a; Extended data Fig. 4a). Moreover, clade IIa strains had higher number of contigs, higher misassemblies and lower genome size than the rest of the strains sequenced for this study (H = 18.82, df =1, *P* <0.001; Extended data Fig. 4b-d). To determine if erroneous assembly affected the clustering of this clade, we performed long read sequencing (Pacific Biosciences Inc.) on one strain from this clade and used it in a hybrid approach with Illumina short reads. This resulted in a partially complete chromosome (longest contig - 4.7 Mbp) but 30 non-circular contigs were not scaffolded. An alignment of the ordered contigs with the ancestor showed that the strain did not suffer significant gene loss as suggested through short or long reads only assembly (Supplementary Table 1), however a 3.2 Mbp chromosomal region was inverted (Fig. 3b). This strain also carried an untypeable circular episome that carried uncharacterized proteins and two genes encoding NADH-quinone oxidoreductase (*nuo*) (∼22 kbp) (Extended data Fig.4e), and an IncX1 plasmid with divergent protein sequences from the IncX1 present in the ancestor (Extended data Fig.4f). Gene inversions can increase the diversity and virulence of pathogens^19^, hence we questioned if chromosomal inversion was present in other evolved strains. To do this, we aligned protein sequences of two complete circular chromosome from trial 1 and 2 with the ancestor. This analysis showed that the isolate from trial 1 harbored a ∼2 Mbp chromosomal inversion (Extended data Fig. 5a).

**Figure 4.**
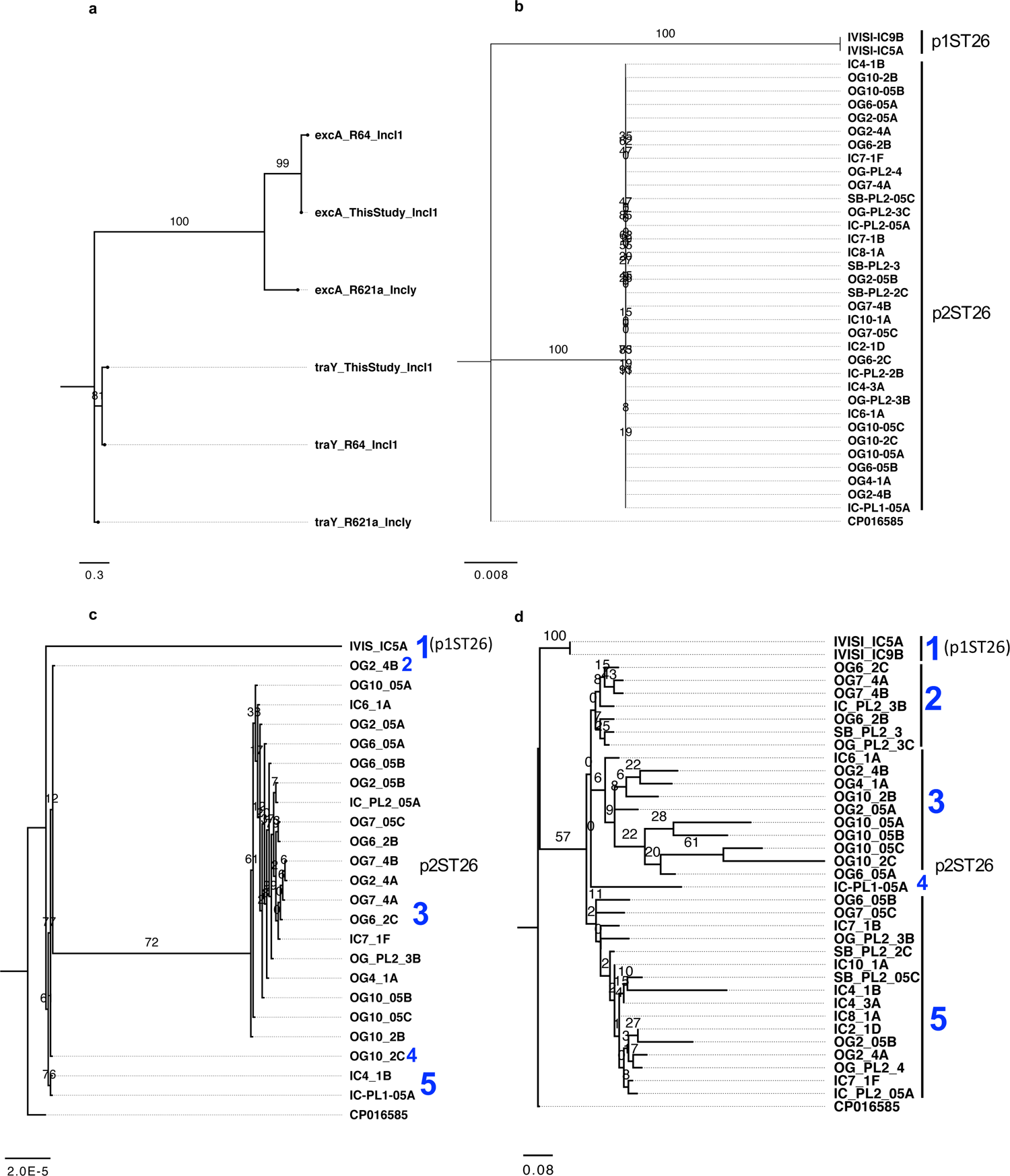
Gene flux contributed to the diversity of IncI1-pMLST26 plasmids. **a,** Maximum likelihood tree constructed using TraY and ExcA protein sequences from a representative IncI1 plasmid from this study, R64 (IncI1) and R621a (IncI1-gamma). The GTR model of nucleotide substitution and the GAMMA model of rate heterogeneity were used for sequence evolution prediction. Tree was rooted with traY sequence of R621a. **b - d,** Maximum likelihood tree of pST26 IncI1 plasmids from this study (n = 36) constructed using complete plasmid DNA sequences, core genes (**c**) and accessory genes (**d**). The GTR+I, GTR and JC+I model of nucleotide substitution and the GAMMA model of rate heterogeneity were used for sequence evolution prediction for b, c, and d, respectively (Tree was rooted with the closest relative (GenBank: CP016585) found through NCBI BLASTn)). Clade numbering were arbitrary assigned to show number of clades found and not order of evolution. Numbers shown next to the branches represent the percentage of replicate trees where associated isolates cluster together based on ∼100 bootstrap replicates.

**Figure 5.**
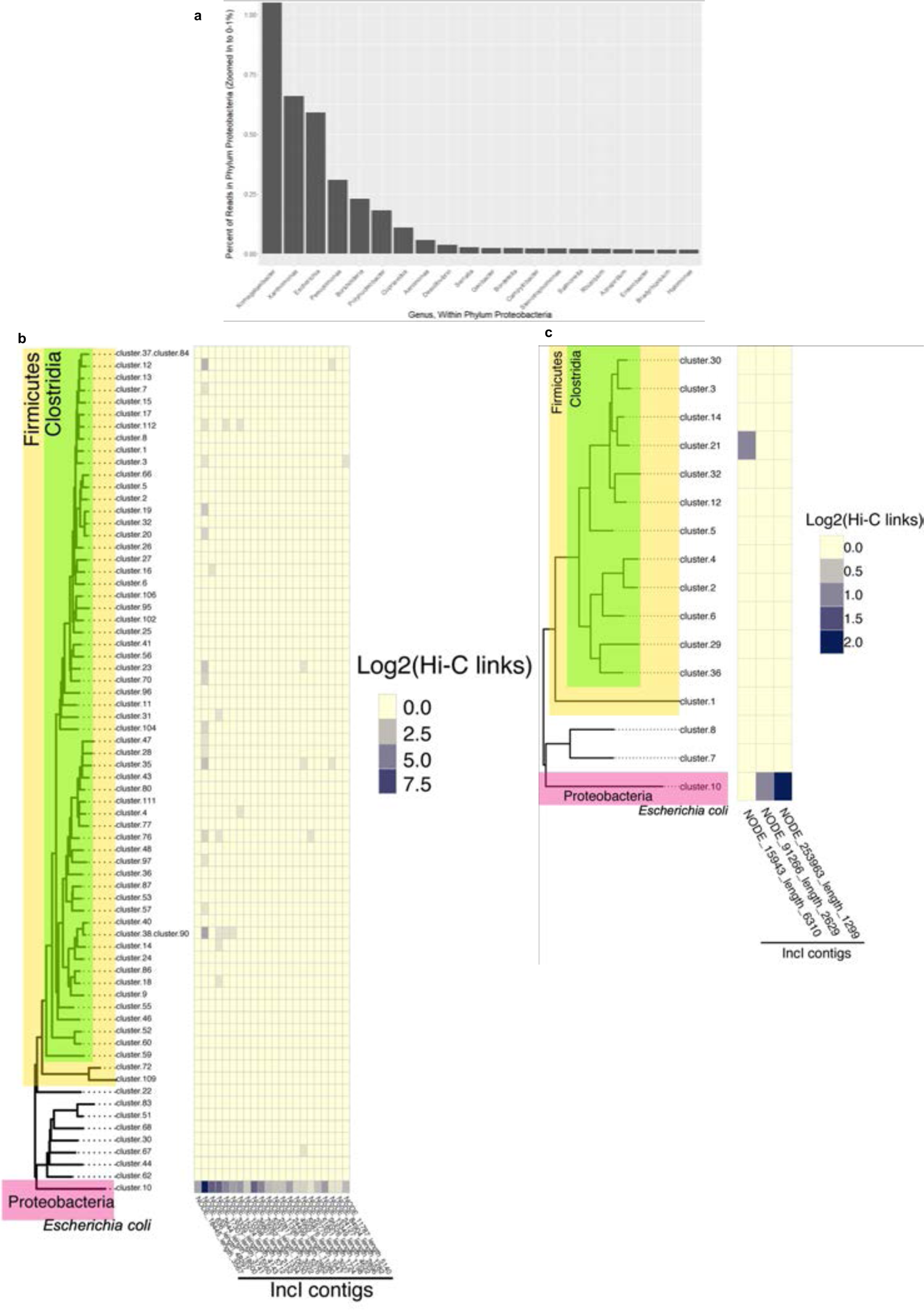
*Proteobacteria* was the reservoir of IncI1-plasmids in broiler chickens. **a**, Percentage of metagenomic reads assigned to the top 20 bacterial genera of the phylum Proteobacteria present in samples Hi-IC-FL1 and Hi-IC-FL2. Only reads assigned to phylum *Proteobacteria* (70.9 % of the metagenomic reads) were used and zoomed in to show 0-1% of total reads within phylum Proteobacteria (96.7% of reads within phylum *Proteobacteria* were assigned to *Komagataeibacter*). **b and c,** IncI1 contigs hosts found by Hi-C contacts. IncI1-derived contigs (horizontal axis of heatmap) show specific Hi-C associations with metagenome-assembled genomes (MAGs) present in samples Hi-IC-FL1 (**b**) and Hi-IC-FL2 (**c**) (vertical axis of heatmap). MAGs are derived from Hi-C deconvolution of the metagenome assembly and placed into a bacterial phylogeny using CheckM. Cluster.10 in each sample is an *Escherichia coli* genome. In Hi-IC-FL1, one MAG (cluster.82) representing an extremely fragmented and partially contaminated *Escherichia coli* genome is omitted for clarity (due to its contamination, this cluster was placed in *Clostridia* by CheckM). Heatmap values indicate transformed counts of Hi-C read contacts (indicating intracellular physical proximity of IncI1 contigs to those genomes. Heatmap values were pseudocounted to facilitate plotting of log-transformed data including zeroes.

Gene inversion changes the leading or lagging strand sequence to its reverse complement, thus altering the GC skew of the affected gene or genome from a positive value to a negative value or negative to a positive^19^. Accordingly, we confirmed that there was a GC skew inversion in these regions (Extended data Fig. 5b and c). Gene inversions occur after rejoining of DNA breaks and inverted genes can introduce sequencing bias and misassembly^20^. Here, deletions affecting the cytochrome c maturation gene cluster (*ccm*) contributed to the inversion of genes in SH strain *og8-05a* (Fig. 3b; Extended data Fig. 5b). Further, the *oriC* of *og8-05a* was reoriented when compared to the ancestor. On the other hand, deletion of a putative YebC/PmpR transcriptional regulator and a deletion between *cysK* and *cysZ* led to the reorientation of *oriC* in strain *ic9b* (Extended data Fig. 5a and c). The inverted region in these genomes encoded 34 - 60 % of the 168 virulence genes present in the ancestor including *Salmonella* pathogenicity islands 1 and 2, fimbrial and adherence genes, type 1 and 2 secretion system proteins and prophages (Supplementary data 2; Extended data Fig. 6). Taken together, this result suggested that gene flux and homologous recombination shaped the genome of SH strains after entry into the broiler chick gut.

**Figure 6.**
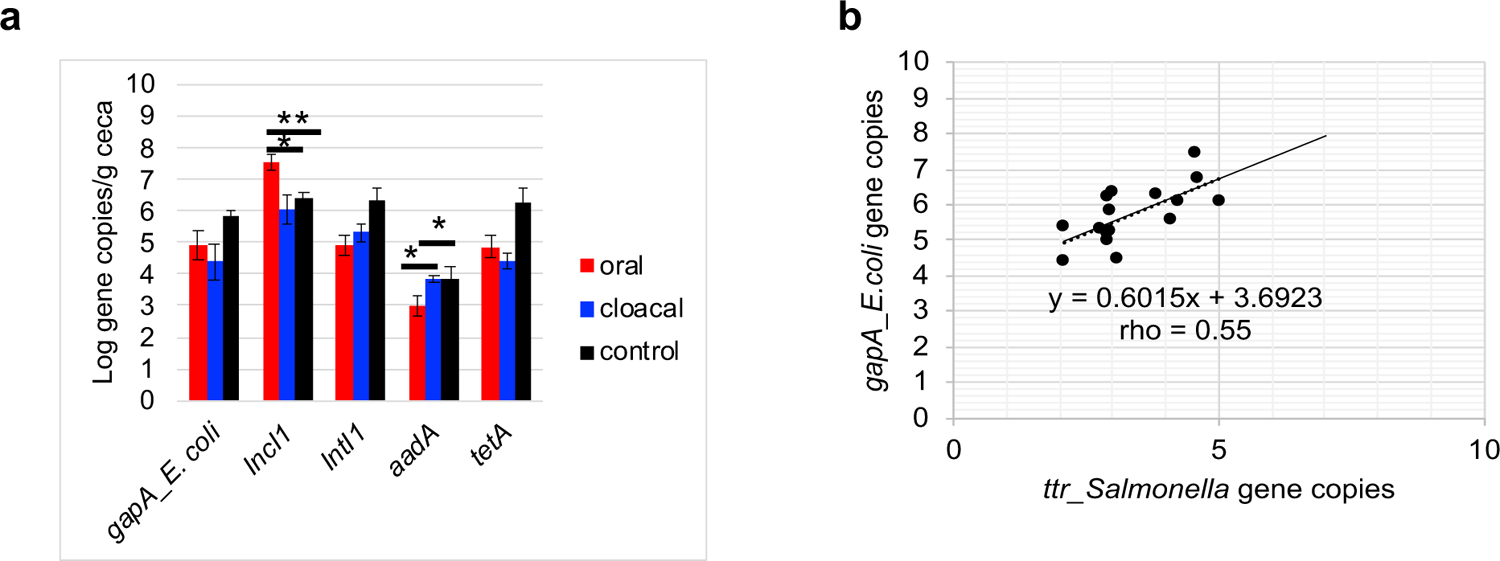
SH challenge affected the relative abundance of antibiotic resistance genes present in the ceca and litter of broiler chickens. **a**, Concentration of selected antibiotic resistance determinants, IncI1 and *E. coli* present in the ceca of broiler chickens. Gene copies were determined using qPCR. Primers targeting the housekeeping gene (*gapA*) and *inc*RNAI was used for *E. coli* and IncI1 quantification, respectively. b, Pearson correlation between *Salmonella ttr* and *E. coli gapA* gene copies present in ceca and litter (n =15).

### pST26 differed in their AR genetic context and genome architecture

To compare pST26 plasmids, we used hybrid assemblies to achieve circular plasmids for two MDR SH isolates recovered from trial 1 and trial 2. The pST26 plasmids from trial 1 (p1ST26) and trial 2 (p2ST26) were determined to be ∼ 112 kbp long and ∼ 87 % identical (Fig. 1f). Both carried the atypical IncI1 backbone including regions encoding replication, stability, leading and conjugative transfer^21^. To determine the IncI1-complex group, we used the classical *traY* and *excA* protein sequences of plasmid R64 (IncI1) and R621a (IncI1-gamma). A reconstructed tree confirmed that both plasmids belong to the IncI1 group (Fig. 4a). A 19.8 kb variable region encoding transposons and AR was found in both plasmids. For p1ST26, this region encoded *aadA1, aac(3)-Via*), *sul1* and quaternary (quats) ammonium compound resistance *(qacEΔ1*) (Fig.1f; Extended data Fig. 7). In contrast, this region carried *tetA* and *mer* operon in addition to *aadA1* and *aac(3)-Via* in p2ST26 (Fig. 1f).

**Figure 7.**
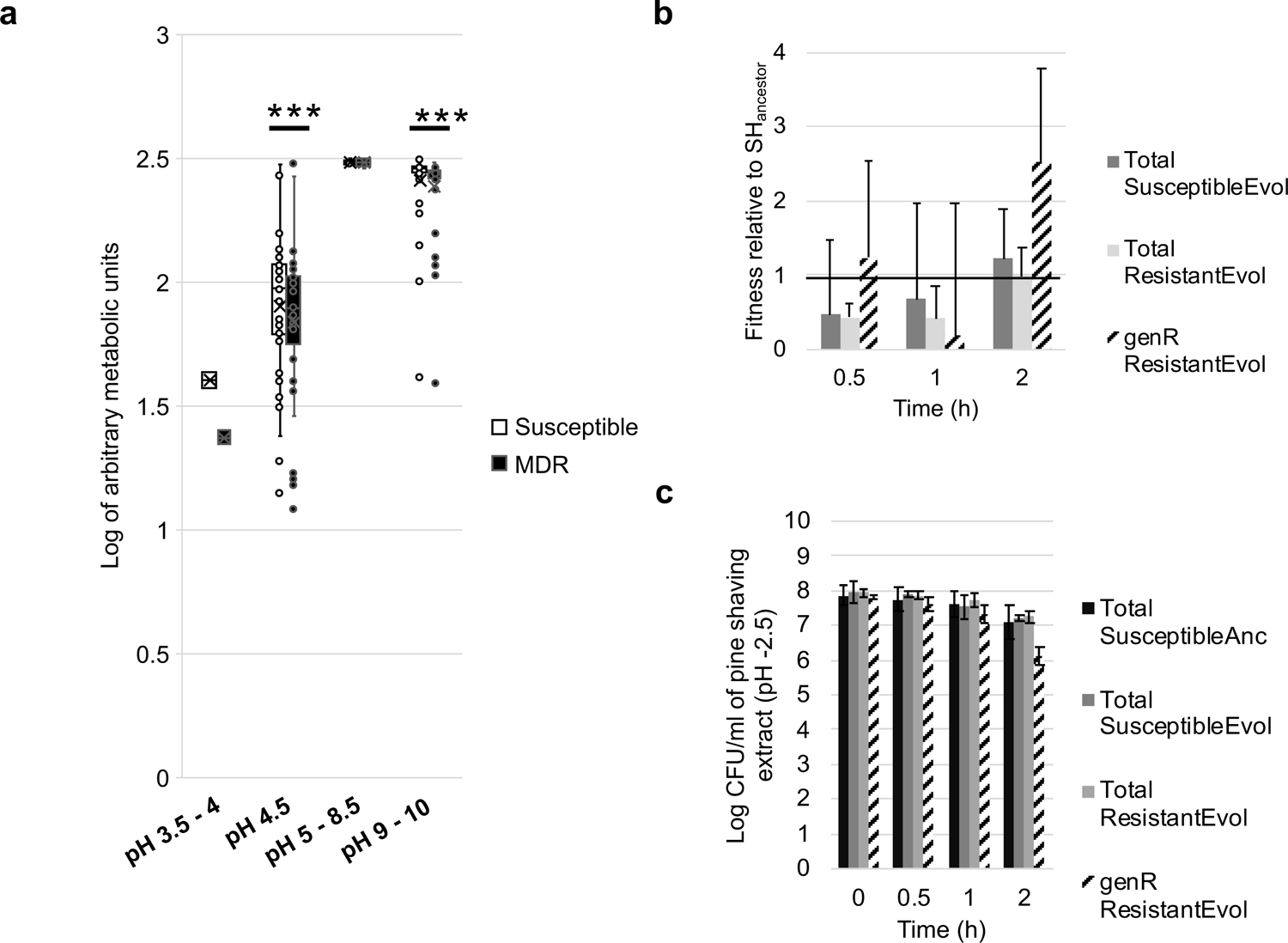
SH exposure to acidic pH imposes negative frequency dependent selection for IncI1 (pST26). **a** Box plot comparing the metabolic activity of one evolved susceptible (no IncI1) and one multidrug (MDR carrying pST26) SH strain using phenotype microarray (PM) plates (Biolog Inc). Pre-configured PM plates are composed of microtiter plates with one negative control well and 95 wells pre-filled with or without nitrogen and at pH of 3.5 – 10 (n = 1 plate per strain; no of wells per plate for pH 3.5-4, 4.5, 5-8.5 and 9-10 was 2, 6, 37, and 38 respectively; \*\*\**P* < 0.001 (Wilcoxon signed rank test). **b and c,** Fitness and abundance of evolved susceptible and MDR SH strains when exposed to pine shaving extract of pH 2.5. Each bar represents the average fitness or abundance (colony forming units per ml of pine shaving extract) of three individual population that was established from three single bacterial colonies from three different strains (Error bar = Standard deviation). The horizontal line in **b** represents the fitness of three SH-Ancestor population established from three single bacterial colonies. (Anc = Ancestor, Evol – Evolved i.e., SH isolate recovered during *in vivo* experiment, genR – population resistant to gentamicin).

To investigate whether gene flux influenced IncI1 genome architecture, we used p1ST26 and p2ST26 as reference genomes for the rest of the MDR isolates carrying IncI1. By aligning raw reads to complete pST26 genomes, we were able generate a consensus IncI1 plasmid contig for each MDR isolate. A tree built with these plasmids resulted in two clades represented by p1ST26 and p2ST26 plasmids, respectively (Fig. 4b). Multiple alignment of protein sequences revealed that pST26 plasmids differed in number of genes carried and gene alleles. This was pronounced for regions encoding AR, pilus/shufflon assembly proteins (*pil*) and IS66-family of transposases (Supplementary data 3). A pangenome analysis revealed that the pST26 from this study carried 70 core genes (genes present in ≥ 95% of the plasmids) and 90 accessory genes (genes present in < 95% of the plasmids). A tree reconstructed with the core genes and accessory genes divided the pST26 into 5 major groups (Fig. 4d and c). For both trees, the p1ST26 formed one clade and the p2ST26 made up the other clades. No SNPs were found between p2ST26 but p1ST26 differed from p2ST26 with 21 SNPs and 2 insertions (Supplementary data 3; Supplementary Table 2). The insertions were present in only one p1ST26 plasmid. One insertion was found on *tniA* (a DDE-type integrase/transposase/recombinase protein) and the other was between tn3-like transposon and *yadA*. Inserted DNA showed significant homology to *tniA* present in *E. coli* plasmid EcPF5 (Genbank: CP054237) and IS5057 present in *E. coli* plasmid pIOMTU792 (Genbank: C542972), respectively.

### *E. coli* was the major IncI1 reservoir in broiler chicks

The application of proximity-ligation method (Hi-C) has improved the assembly of metagenomes and made it possible to detect plasmid-host associations^22^. Therefore, we used Hi-C to determine the bacterial species carrying IncI1 plasmids in the broiler chicken microbiota. Hi-C was done on two cecal samples from chicks challenged with SH via the cloaca (hereafter referred to as Hi-IC-FL1 and Hi-IC-FL2). We chose IC samples since all chicks in this group were successfully colonized with SH compared to OG or SB. Our goal was to determine if Hi-C could find the bacterial host of IncI1 plasmids and detect inoculated SH strains. *Salmonella enterica* was found in Hi-IC-FL1 and Hi-IC-FL2 using metagenome sequence classification bioinformatic tools^23, 24^ (Fig. 5a). However, serovar determination with cluster contigs (∼ 1.4 and 1.2 Mbp total contig size, respectively) was not possible with *Salmonella* serovar prediction tool^25^ or after alignment of contigs to SH reference genome. We detected IncI1 plasmid contigs in Hi-IC-FL1 that were linked to *E. coli* clusters (Fig. 5b and c; Supplementary Table 3). In addition, we found contigs (n = 151, mean contig size = 17,607 bp, smallest contig = 1,027 bp, largest contig = 282,389 bp) with protein sequences identical to pST26 in both Hi-IC-FL1 and Hi-IC-FL2 (Supplementary data 4). Other bacterial clusters belonging to the *Firmicutes* showed a low level of linkage to IncI1 (∼100X less in Hi-IC-FL1). These results suggested that *E. coli* was the main reservoir of IncI1 plasmids.

Next, we investigated if SH challenge affected *E. coli* and IncI1 plasmid abundance. To answer this, we used qPCR primers targeting the glyceraldehyde-3-phosphate dehydrogenase (*gapA*) gene of *E. coli*, the *inc*RNAi region of IncI1 and selected genes carried on p2ST26 (*intI1*, *aadA* and *tetA*) to determine their concentration in the ceca and litter. Litter carried higher concentration of *E. coli*, IncI1 and p2ST26 genes than ceca on a per gram basis (W = 1, *P*< 0.001). (Fig. 6a; Extended data Fig.8a). In the ceca, IncI1 was higher in OG compared to IC or CTRL (X^2^ = 9.17, df =2, *P* = 0.01), while *aadA* (H = 8.25, df = 2, *P* = 0.016) was lower in OG compared to IC or CTRL (Fig. 6a). CTRL chicks carried a higher concentration of *E. coli* (X^2^ = 2.32, df =2, *P* = 0.31), *tetA* (X^2^ = 5.74, df =2, *P* = 0.05) and *intI1* (X^2^ = 5.69, df =2, *P* = 0.05) genes than OG and IC. A principal component analysis with these genes and *Salmonella* tetrathionate reductase (*ttr*) revealed that samples within the control and challenge groups clustered with their respective groups and that the challenge and control groups were separated. (Extended data Fig. 8b). A correlation analysis between *Salmonella* and *E. coli* for challenged chicks showed a significant positive correlation (S = 784, ρ = 0.55, *P* =0.007) (Fig. 6b). These results suggested that the presence of SH affected *E. coli*, IncI1 and ARG concentrations.

### Broiler chicks carried *E. coli* strains harboring pST26 IncI1 plasmids

To confirm if broiler chicks from this study harbored *E. coli* populations that could serve as a reservoir of pST26 plasmids, we retrospectively screened the cecal contents of the broiler chicks used for Hi-C on CHROMagar™ supplemented with gentamicin and tetracycline (see supplementary methods) and selected five colonies for whole genome sequencing. As expected, we found pST26 in two strains (phylogroup F, MLST 6858), while the other three strains (phylogroup D, MLST 69) carried an IncI1 that harbored no ARG, and an unknown MLST (Table 1). Next, we compared the pST26 in SH to that found in *E. coli* using complete plasmids. The p1ST26 and p2ST6 IncI1 plasmids of SH were determined to be 86.8% and 99.4 % identical to pST26 present in *E. coli*. The main difference was is in the region encoding AR and transposases in p1ST26 (Extended data Fig 9).

**Table 1:** Multidrug resistant E. coli genomes found in the ceca of broiler chicks

| Isolate ID | Resistance phenotype<sup>†</sup> | Antibiotic resistance genes<sup>‡</sup> | Plasmids<sup>§</sup> | Phylogroup | MLST |
| --- | --- | --- | --- | --- | --- |
| Ec-FL1-1X | **Amp Gen**<br>**Str Fis Tet** | ***aac(3)-lid aph(3')-la***<br>***aph(6)-ld aph(3'')-lb***<br>***blaTEM-1B tet (B) sul2*** | IncF:B:A-, IncI1 | D | 69 |
| Ec-FL2-2X | **Amp Gen**<br>**Str Fis Tet** | ***aac(3)-lid aph(3')-la***<br>***aph(6)-ld aph(3'')-lb***<br>***blaTEM-1B tet (B) sul2*** | IncF:B:A-, IncI1 | D | 69 |
| Ec-FL2-5X | **Amp Gen**<br>**Str Fis Tet** | ***aac(3)-lid aph(3')-la***<br>***aph(6)-ld aph(3'')-lb***<br>***blaTEM-1B tet (B) sul2*** | IncF:B:A-, IncI1 | D | 69 |
| Ec-FL1-2X* | Gen Str Tet | *aadA1 aac(3)-Via tet(A)* | IncF:B:A- IncI1 IncI2<br>Col8282 ColRNAI | F | 6858 |
| Ec-FL1-5X* | Gen Str Tet | *aadA1 aac(3)-Via tet(A)* | IncF:B:A- IncI1 IncI2<br>Col8282 ColRNAI | F | 6858 |

\* *E. coli* isolates carrying IncI1-pST26 with >99% identical DNA sequence with p2ST26 acquired by SH.
<sup>†</sup> Amp – ampicillin, Gen – gentamicin, Str – streptomycin, Tet – tetracycline, Fis – sulfisoxazole.
<sup>‡</sup> Boldness denotes antibiotic resistance genes encoded on the chromosome.
<sup>§</sup> Illumina short reads and PacBio long reads were combined to assemble the plasmids of *E. coli* strains Ec-FL1-1X and Ec-FL1-2X.

### pST26 carriage did not pose a fitness cost under selection pressure

pST26 carried accessory genes for AR, metal resistance or disinfectants (i.e., quaternary ammonium compounds) Furthermore, SH strains carrying pST26 differed by route of exposure e.g., higher proportion in oral vs. cloacal. To this end, we questioned if any of these factors could exert a selective pressure for pST26. We did not administer antibiotics to broiler chicks throughout study, therefore we did not consider antibiotic selective pressure. However, we detected metals including nickel, chromium, copper, and zinc in the starter feed in parts per million and we found contaminants such as arsenic, lead and cadmium in trace amounts (Supplementary Table 4). Disinfectants including quaternary ammonium compounds were used for the cleaning of broiler houses and equipment, thus we considered disinfectant as another source of selective pressure in this study. To determine their effect on SH fitness, we used phenotype MicroArray™ (PM) 96-well plates supplemented with selected metal chlorides and disinfectants at varying concentrations to compare a pST26 carrying strain to a susceptible evolved strain. There was no significant difference in metabolic activity for SH strain carrying pST26 compared to its susceptible counterpart for metals and disinfectant but pST26 exhibited higher metabolism for benzethonium chloride, chromate, tellurite, and zinc, whereas the susceptible stain showed higher metabolism for dequalinum chloride, chromium, and cesium (X^2^= 1.83, df = 1, *P* = 0.1762) (Extended data Fig. 10a and b).

Cox and colleagues have suggested that the acidic pH of the upper GI tract was associated with the lower number of chicks positive for *Salmonella* and *Campylobacter* after oral gavage compared to cloacal inoculation^26–28^. Based on this premise, we hypothesized that SH strains carrying pST26 will survive better under exposure to acidic pH. To answer this question, we first compared the survival of the strains under different pH levels (3.5 - 10) with or without nitrogen sources and amino acids using PM plates. SH strain carrying pST26 exhibited lower metabolic activity compared to the susceptible strain at pH 3.5 – 4 (V = 3; *P* = 0.5), pH 4.5 with and without nitrogen sources (V = 599; *P* = 3.426e-06) and at pH 9.5-10 with and without sources of nitrogen (V = 714; *P* = 6.547e-07) (Fig. 7a). Next, we compared the survival of the susceptible ancestor, susceptible evolved strains without pST26 and antibiotic resistant evolved strains carrying pST26 in pine shaving extract (PSE) adjusted to pH – 2.5. We acclimatized the bacterial population to PSE (pH – 6.5) for 2 h before exposure to acidified PSE (Extended data Fig. 10c). We screened for pST26 carrying populations using the gen^R^ marker on pST26. After 2h of exposure to acidified PSE, the SH population carrying pST26 (resistant plus susceptible cells) did not suffer a fitness cost, and cells carrying pST26 exhibited higher fitness than their susceptible counterparts (X^2^= 4.35, df = 2, *P* = 0.113) (Fig. 7b). Notwithstanding, pST26 SH cells had a lower population size than the ancestor and evolved susceptible strains (X^2^ = 6.37, df =3, *P* = 0.095) (Fig. 7c). This result suggested that pST26 carriage does not pose a fitness cost on SH host but benefits the host when exposed to acidic pH.

## Discussion

In this study, neonatal Cobb 500 broiler chicks challenged with a susceptible *S*. Heidelberg (SH) strain carried susceptible and multidrug resistance SH population 2-weeks after in their ceca and litter. This resistance phenotype was conferred by the acquisition of IncI1-pST26 plasmids. pST26 plasmids are present in *S*. Heidelberg, *S*. Typhimurium, *S*. Anatum, *S*. Derby, *S*. Schwarzengrund, *S*. Saintpaul and *E. coli* strains isolated from animal and human sources (Extended data Fig. 11). The closest IncI1 plasmids to pST26 from this study are pST26 present in *E. coli* (CP018625) and *S*. Typhimurium (CP027409). The plasmids carry aminoglycoside resistance genes with or without tetracycline, sulphonamide, *qac* and *mer* resistance genes, making it a multidrug/disinfectant/metal resistance plasmid. Additionally, pST26 transfer was higher in chicks challenged orally compared to chicks inoculated via the cloaca. This route specific rate of conjugation suggested that the selection for pST26 in our study was associated with the differential pH in the upper GI versus lower GI tract. This was corroborated by in vitro experiments, where acidic pH reduced the metabolism of SH cells carrying pST26 compared to cells with no pST26. Furthermore, carriage of pST26 increased the fitness of pST26-carrying cells compared to the susceptible SH population after exposure to acidic pine shaving extract. This type of acid-imposed selection for IncI1 alludes to the “negative frequency dependent selection hypothesis” - where the fitness of the population is higher when a plasmid is present at a low frequency in the population^29^.

This could also explain the higher levels of SH, but lower positivity rate found in the ceca of broiler chicks gavaged compared to cloacally challenged chicks^16^ (Fig. 1b). There are multiple barriers that makes it challenging for SH to colonize the ceca of broiler chicks gavaged including the acidity of the upper GI tract and the length of the GI tract to be traveled. When we measured the distance of the cloaca and esophagus to the ceca of a 49-day old Cobb-500 broiler chicken (photo not shown), we estimated that the SH inocula will have to travel ∼20X farther to get to the ceca of gavaged chicks compared to cloacal inoculated chicks. It is plausible that this longer “residence” time in the upper GI allows more opportunities for SH to make contact and compete with other members of chicken microbiota. In this study, such an event gave rise to SH cells that acquired pST26 from *E. coli*. It is also noteworthy that AR acquisition occurred at a lower rate during trial 1 compared to trial 2. This could be due to several factors that warrants further study - including the higher pH and moisture of the litter in trial 1 compared 2, seasonal differences (trial 1 was done in the fall and trial 2 was in the spring) and the antibiotic/disinfectant practices in place at the hatchery or breeder farms at time the experiments were conducted.

The IncI1-complex are widespread in *Enterobacteriaceae* and have been reported to be the major driver of AR in *Salmonella* and *E. coli*. Our understanding of their mode of transfer and AR evolution have been limited to studies on their prevalence, mating experiments and in vivo challenge with mice^21, 30, 31^. Fischer et al^32^ showed that *E. coli* populations in broiler chickens acquired β-lactam resistance after 4-days old broiler chicks were gavaged with an *E. coli* strain carrying *bla_CTX-M-1_* on an IncI1 plasmid. In a twin pair experiment, Hagbo et al^33^ revealed that IncI1 carrying *bla_CMY-2_*, ColE1 and P1 bacteriophage are frequently transferred between microbiota present in pre-term infant gut. Recently, Nyirabahizi,et al^34^ used likelihood-ratios to show that *E. coli* AR predicted *Salmonella* resistance to β-lactams in retail meat, and to gentamicin, ceftriaxone, and amoxicillin-clavulanic acid in cecal samples and suggested that the horizontal transfer of *bla_CMY-2_* played a role. These studies have demonstrated that IncI1 is the most prevalent plasmid carrying β-lactam in *Enterobacteriaceae* and that they are transferable from *Salmonella* to *E. coli* and vice-versa. IncI1 plasmids can harbor transferable gene cassettes with AR variants for β-lactam (*bla_TEM_*, *bla_CMY_*, *bla_CTX_*), aminoglycoside (*aadA*, *aacA*, *aph*), tetracycline (*tet*), sulphonamide (*sul*), fosfomycin (*fosA*), phenicol (*cmlA*, *floR*) and trimethoprim (*dfrA*) resistance. Furthermore, mobile elements could carry AR genes for disinfectants and metals including ammonium compounds (*qac, sug*), mercury (*mer* operon) and arsenic (*ars* operon).

The nutrient-niche hypothesis suggests that SH will only be able to invade a new niche if it can metabolize a growth-limiting resource or if it can outcompete a resident species and a vacant niche arises due to a new component in diet or by eliminating the competitor ^35^. Competition for oxygen was shown to be the limiting factor that contributed to the virulence of *S*. Enteritidis towards *E. coli* in neonatal chickens^36^. Competition within and between SH and *E. coli* strains played a key role selecting for antibiotic resistant SH in our study. SH challenge perturbed the bacterial community and ARGs of broiler chicks and SH population was positively correlated with *E. coli*. Consequently, gene flux and genome inversion shaped the evolution of SH in broiler chicks. For SH and *E. coli* to co-evolve and persist in the same new niche, recombination of genes occurred and led to the emergence of new SH strains. In our study, pST26 emerged after conjugation between *E. coli* donors from broiler chicks raised on fresh litter and SH. As expected, SH genomes were grouped into nine core genome MLST clusters with eight clusters representing new strains (Extended data Fig. 12). These species/strains are expected to be fitter in the new niche and persist longer.

## Materials and Methods

### SH inoculum preparation

We have previously described the characteristics of the SH strain (SH-2813nal^R^) used for this study and how the inoculum was prepared^16^. Briefly, the strain belongs to the multilocus sequence type 15, carries a 37-kb conjugative plasmid, and is resistant to erythromycin, tylosin, and fosfomycin. The strain was recovered from broiler chicken carcass in 2013 and made resistant to 200 ppm of nalidixic acid (nal^R^) for selective enumeration. The nal^R^ phenotype is conferred by a serine to tyrosine substitution at position 83 of DNA gyrase subunit A protein (GyrA). The SH inoculum was grown overnight in poultry litter extract, centrifuged, and resuspended in 1X phosphate buffered saline (PBS). The resuspended cells were used as inocula. The genome of the ancestor to SH-2813nal^R^ was sequenced using Illumina and PacBio sequencing to achieve a complete circular chromosome and plasmid (Genbank). SH-2813nal^R^ carries six mutations (Supplementary data 5) including the mutation affecting *gyrA* that are **not present in the ancestor.**

### Challenging broiler chicks with SH

One-day-old Cobb 500 broiler chicks were purchased from a commercial hatchery in Cleveland, GA, USA. Upon purchase, chicks were placed in plastic crates lined with brown paper and transported to the University of Georgia, Poultry Research Center (33.90693362620281, −83.37871698746522). One hundred chicks were either uninoculated, gavaged or cloacally inoculated with 100 µl-volume of SH inoculum (Fig.1a). We included a seeder-bird (SB) colonization method, whereby five orally gavaged chicks were mingled with twenty uninoculated chicks. Each inoculated chick received ∼10^6^ colony forming unites (CFU) of SH-2813nal^R^. Afterwards, chicks were placed in floor pens at a stocking density of 0.65 m^2^/chick on fresh pine shavings litter. Broiler chicks were given water and feed ad libitum and raised antibiotic-free on starter diet for 2-weeks (starter feed was synthesized by the University of Georgia’s poultry research center’s feed mill). The concentration of metals and priority pollutants in feed was determined by the University of Georgia’s feed and environmental water laboratory (Athens, GA, USA) (Supplementary Table 4). Husbandry and management followed commercial broiler chicken industry guidelines. After 2-weeks, ten chicks from OG, IC and control groups and fifteen chickens from SB (five seeder chicks and ten uninoculated pen mate) were sacrificed to determine the extent of SH colonization in ceca. The experiments were done in two trials conducted in September 2017 (Trial 1) and April 2018 (Trial 2). The study was approved by the University of Georgia Office of Animal Care and Use under Animal Use Protocol: A2017 04-028-A2.

### Cecal and litter bacteriological analyses

Ceca were aseptically removed from the eviscera of 2-week-old broiler chicks, placed in a stomacher bag and transported on ice to the US National Poultry Research Center for analysis. Ceca were weighed and buffered peptone water (BPW) (BD Difco, MD, USA) was added 3X volume to the weight (v/w) and stomached for 60s. Serial dilutions were made and plated onto Brilliant Green Sulfa agar (BGS) (BD Difco, MD, USA) containing 200-ppm nal. In addition, a 10 µl inoculating loop was used to streak cecal slurry (cecal contents in BPW) onto Xylose Lysine Tergitol-4 agar (BD Difco, Sparks, MD). supplemented with 4 ppm tetracycline for trial 1 and BGS supplemented 32 ppm ampicillin (amp) or 32 ppm streptomycin for trial 2. All antibiotics were purchased from Sigma-Aldrich (St. Louis, MO), unless otherwise noted. Plates were incubated along with the cecal slurry for 24 h. All bacterial incubations were carried out at 37°C, unless otherwise noted. After incubation, colonies were manually counted and serial dilution plates with 2 – 100 colonies were used for CFU per gram calculation. If no colonies appeared on serial dilution plates, cecal slurry was streaked onto a new BGS plus nal plate and incubated overnight. After incubation, plates were examined for the presence/absence of *Salmonella* colonies.

Broiler chicken litter was collected as grab samples from seven locations (4 corners of the pen and under the waterer) in each pen after chicks were removed. The litter samples were pooled and 30g was processed in duplicates from each pen as previously described^37^. Serial dilutions of the litter slurry were made and plated onto BGS agar containing 200 ppm nal. Plates were incubated overnight, and colonies were manually counted and reported per gram litter dry weight. Litter pH and moisture was determined as described previously^37^. We selected randomly 2 – 6 single colonies from BGS plates supplemented with nal from each ceca and litter sample and archived them in 30% LB glycerol at −80 °C. In addition, cecal slurry was saved at a 4:1 ratio in Luria Bertani broth (BD Difco, MD, USA) containing 30% glycerol at –80°C, whereas litter samples were stored in vacuum sealed whirl pak bags at −20°C.

### Determining SH metabolism under selective pressure

To determine if exposure to metals, disinfectants or acidic pH poses a selection on SH strains carrying IncI1 plasmids, we used 96-well Phenotype Microarray (PM) ™ MicroPlates (PM10, PM12B, PM13B, PM15B and PM16A) (Biolog, Inc., Hayward CA) to compare the metabolic profile of one gentamicin resistant strain harboring IncI1 and one susceptible strain. PM™ plates uses cell respiration via NADH production to determine cell metabolic activity. If the phenotype is positive in a well, the cells respire actively, reducing a tetrazolium dye and forming a strong color (Biolog, Inc.). Briefly, SH strains were sub-cultured twice in universal growth agar (BUG+B) for 24 h at 37°C. Cells were removed with a sterile swab and transferred to 16 ml IF-0 to achieve a final turbidimetric transmission of 42%T. Afterwards, 15 ml of the 42%T cell suspension was transferred to 75 ml of IF-0 +dye to achieve a final transmission of 85% T before 600 μl of the cell suspension was transferred to 120 ml IF-10+ dye. The suspension was mixed and each well in a microplate was inoculated with 100 μl of the suspension. After inoculation, microplates were covered with a sterile plastic film and monitored automatically for color development every 15 min for 48 h at 37 °C using and OmniLog reader for 48 h. To identify phenotypes, the kinetic curves of the gentamicin resistant and susceptible strains were compared using OmniLog PM software. For each PM plate, the respiratory unit for each well at 22 h was extracted and log_10_ transformed. All supplies used were purchased from Biolog, Inc.

### Exposing SH to acidic pH

To determine the fitness of evolved SH strains that acquired antibiotic resistance compared to evolved susceptible strains after exposure to acidic pH, we exposed gentamicin resistant (n = 3) and susceptible (n =3) strains recovered from the litter of chicks gavaged with SH to acidified filter sterilized pine shaving extract. Pine shaving extract (PSE) was prepared as described for poultry litter extract^37^, using fresh pine shavings collected from the University of Georgia, Poultry Research Center, Athens, GA, USA. Three single colonies of each strain including the ancestral SH-2813nal^R^ were selected from overnight cultures grown on sheep blood agar and transferred to a microcentrifuge tube containing 900 μl of PSE (pH = 6.52) i.e., one tube per strain. After transfer, tubes were vortexed, covered with a gas permeable paper strip and incubated at 41°C under microaerophilic conditions (5% O_2_, 10% CO_2_, and 85% N_2_) for 2 h. After incubation, tubes were vortexed and 100 μl of the suspension was transferred to a microcentrifuge tube containing 900 μl of PSE (pH of PSE was adjusted to 2.54 using 1M HCl (Spectrum Chemical Mfg. Corp., CA, USA) and 1M NaOH (Fisher Chemical, NJ, USA) Afterwards, tubes were vortexed, and incubation continued for another 2 h as was done for PSE at pH 6.5. To determine SH population in PSE, one replicate tube per strain was removed and serially diluted in 1X PBS at timepoints 0 and 2 h for pH 6.5 and 0.5, 1 and 2 h for pH 2.5. Serial dilutions were plated onto BGS agar plates supplemented with or without 16 ppm gentamicin. SH colonies were counted 18 - 24 h after incubation using a calibrated automated colony counter.

The fitness of each evolved SH population relative to the SH-2813nal^R^ population was determined after 2h at pH 2.54 as described by San Millan et al^38^:

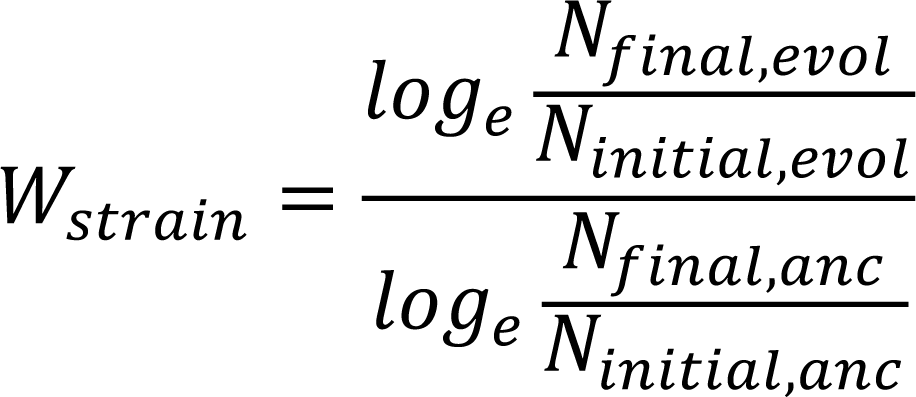

where W_strain_ is the fitness of the evolved susceptible (total susceptible) or gentamicin resistant populations (total resistant and gen^R^ resistant), N_initial, evol_ and N_final,evol_ are the numbers of cells (in CFU) of the evolved susceptible or gentamicin resistant population at timepoint 0 and 2 h after exposure to PSE, and N_initial,anc_ and N_final,anc_ are the numbers of cells of SH-2813nal^R^ population at timepoint 0 and 2 h.

### Antimicrobial susceptibility testing

We performed antimicrobial susceptibility testing (AST) on two hundred and fifty SH isolates recovered from the ceca and litter of broiler chicks following National Antimicrobial Resistance Monitoring System (NARMS) protocol for Gram-Negative bacteria. Minimum inhibitory concentration for isolates were determined by broth microdilution using the Sensititre semiautomated antimicrobial susceptibility system (Thermo Fisher Scientific, Inc., MA, USA) and interpreted according to clinical and laboratory standards institute guidelines when available; otherwise, breakpoints established by NARMS were used. AST was also done on *E. coli* isolates (n = 5) recovered from the ceca of two broiler chicks challenged intracloacally with SH. The Kirby-Bauer disc diffusion assay for tobramycin, kanamycin, neomycin, netilmicin was done on four gentamicin resistant SH isolates as previously described^37^.

### DNA extraction

DNA was extracted and purified from bacterial colonies using FastDNA spin Kit (MP Biomedicals, LLC, CA, USA), whereas 250 mg of cecal and litter were extracted with Qiagen DNeasy Power-Soil DNA kit (Hilden, Germany). The modifications we made to the manufacturers protocol for DNA extraction have been reported ^37, 39^.

### Whole genome sequencing and processing

Illumina short read sequencing was performed on DNA extracted from SH isolates recovered from ceca and litter. In addition, five *E. coli* isolates recovered from two cecal samples were sequenced. Libraries were prepared using either Nextera™ XT or Nextera™ DNA Flex library preparation kits (Illumina, Inc., San Diego, CA) following the manufacturers protocol. Libraries were sequenced on the Illumina MiSeq platform with 150 or 250-bp paired end reads. Additionally, five SH and two *E. coli* isolates were selected for long read sequencing using Sequel II System (PacBio Biosciences Inc.) or MinION (Oxford Nanopore technology). Preparation and sequencing of long read libraries were done by next generation sequencing core centers of University of Georgia and Colorado State University (see supplementary methods for read quality control and demultiplexing).

Genome assembly, resistome characterization, and quality assessment of long reads was done using Reads2Resistome pipeline v.1.1.1^40^. Reads2Resistome performed hybrid assemblies using either Illumina reads and PacBio or Illumina and MinION reads, using both Unicycler^41^ and SPAdes^42^. In addition, hierarchical genome assembly process (HGAP) assembly was performed using PacBio’s SMRT Link v9.0 analysis software suite using default settings. Assembly quality was assessed by QUAST v5.0.2^43^, using default settings) and genome annotation was done using Prokka^44^, Rapid Annotation using Subsystem Technology (RAST)^45^ and BlastKOALA^46^. We confirmed that all SH isolates were *Salmonella enterica* serovar Heidelberg using *Salmonella* In Silico Typing Resource (SISTR)^25^, and used ClermonTyping^47^ and Multi-Locus Sequence Typing (MLST)^48^ for *E. coli* strains. For resistome characterization reads2resistome used ARG-ANNOT^49^, the Comprehensive Antibiotic Resistance Database^50^, MEGARes^51^, AMRFinderPlus^52^, PlasmidFinder^18^, ResFinder^53^ and VirulenceFinder database^54^. For plasmid typing and IncI1 clonal complex determination, we used plasmid MLST^18^. PHAST^55^ was used to identify prophages present in chromosomal contigs and predicted prophage DNA sequence was annotated with RAST and BlastKOALA. ProgressiveMAUVE v. 1.1.1^56^ and MAFFT v. 1.4.0^57^ implemented in Geneious Prime® v 2020.0.1 were used for aligning and comparing sequences. In addition, a pan genome analysis of annotated assemblies was conducted with Roary^58^. Phylogenetic trees based on core genome and accessory genome were reconstructed using the maximum likelihood (ML) method implemented in RAxML-NG v 2.0.0^59^. When computationally possible, the best model of sequence evolution predicted by jModelTest^60^ was used for tree reconstruction, otherwise the GTR + GAMMMA model was implemented. Lastly, we used the bacterial genomes sequenced in the study to create a BLAST database that can be searched for sequences of interest in Geneious Prime®.

Illumina short reads were assembled de novo into contigs using Unicycler v.0.4.7 and characterized with bioinformatic tools described for long reads. Illumina reads were used to determine single nucleotide polymorphisms (SNPs) and indels present in SH isolates. Alignment of raw FASTQ reads to the genome of SH-ancestor was done using Burrows-Wheeler Aligner (BWA)^61^ and SNPs/indels were called using Genome Analysis Toolkit^62^ as described previously^39^. Variant call format (VCF) files of identified SNPs/indels and the Linux/Unix shell script used have been deposited in Dryad Digital Repository: https://doi.org/10.5061/dryad.4tmpg4f8d. SNP based ML tree was reconstructed by converting merged VCF files to PHYLogeny Inference Package format using PGDSpider^63^. Afterwards, isolates with duplicated SNPs/indels were removed, and a ML tree was drawn using the Jukes-Cantor model of nucleotide substitution and GAMMA model of rate heterogeneity. For other bioinformatic analyses performed including pST26-IncI1 consensus determination through FASTQ alignment, see supplementary methods. All raw FASTQ reads for sequenced bacterial genomes are publicly available under NCBI accession numbers:

### Hi-C and cecal metagenome library preparation and analysis

A shotgun and Hi-C DNA library of two cecal samples was created using Nextera™ XT library preparation kit and Phase Genomics (Seattle, WA) ProxiMeta Hi-C Microbiome Kit following the manufacturer’s instructions. Library sequencing was performed by Novogene corporation (Sacramento, CA, USA) on the Illumina HiSeq platform using 150bp paired end reads. Two libraries were sequenced per HiSeq flowcell lane resulting in a total of ∼125 million shotgun reads, and ∼206 million Hi-C reads per sample. Metagenomic FASTQ files were uploaded to the Phase Genomics cloud-based bioinformatics portal for subsequent analysis. Shotgun reads were filtered and trimmed for quality using bbduk^64^ and normalized using bbtools^64^ and then assembled with metaSPAdes^65^ using default options. Hi-C reads were then aligned to the assembly following the Hi-C kit manufacturer’s recommendations (https://phasegenomics.github.io/2019/09/19/hic-alignment-and-qc.html). Briefly, reads were aligned using BWA-MEM with the −5SP and -t 8 options specified, and all other options default. SAMBLASTER^66^ was used to flag PCR duplicates. Alignments were then filtered with samtools^67^ using the -F 2304 filtering flag to remove non-primary and secondary alignments. Lastly, Hi-C read alignments to the assembly were filtered using Matlock (https://github.com/phasegenomics/matlock) with default options (removing all alignments with MAPQ<20, edit distance greater than 5, and read duplicates). Metagenome deconvolution was performed with ProxiMeta^68, 69^, resulting in the creation of putative genome and genome fragment clusters.

Clusters were assessed for quality using CheckM^70^ and assigned preliminary taxonomic classifications with Mash^71^ and Kraken2 (v2.0.8-beta)^23^. To search the metagenome for ARG and plasmids of interest, we used Geneious Prime® to create a BLAST metagenome database. Metagenome contigs with significant homology (E value < 0.05 and at least 1000 bp of aligned sequence) were considered contigs of the respective plasmid or ARG. Hi-C data were then used to link identified sequences to host genomes and genome fragments within ProxiMeta^22^. Phylogenetic visualizations of Hi-C linkages used the placement of clusters in a large prokaryotic phylogeny as estimated by CheckM^70^. Shotgun and Hi-C reads are publicly available under NCBI accession numbers:

### 16S rRNA gene sequence processing and analysis

Cecal and litter DNA were used for bacterial community analysis through the sequencing of the V4 hypervariable region of the 16S rRNA gene of bacterial genomes. Sequencing was done using the paired-end (250 × 2) method on the Illumina MiSeq platform. Raw 16S rRNA gene sequences were processed with dada2 (version 1.14) in the R environment^72^. Briefly, the reads were trimmed where the quality profile dropped below Q30 (250bp for forward and 240bp for reverse reads) and all reads with more than 2 expected errors were removed entirely. The remaining reads were used to learn the error rates and the samples were pooled prior to sample inference. After merging of forward and reverse reads with the default parameters, chimeric sequences were removed with the removeBimeraDenovo command in the consensus method. High quality reads were then classified against the SILVA database version 132^73^. To avoid bias introduced by spurious amplicon sequence variants (ASVs) or from samples with low sequencing depth, any ASVs with less than 5 reads and samples with less than 5000 reads were removed from the dataset before further analysis. Statistical analysis of microbial communities was performed in the R environment using the packages “phyloseq”, “Ampvis2”, “vegan”, and “MaAsLin2”. Alpha diversity indices were calculated with a dataset rarefied to the minimum sample size (8740 sequences). After assessing normal distribution by qqplots, histograms, and the Shapiro-Wilk normality test, the not normal distributed groups were compared using the Wilcoxon Signed Rank test. Beta diversity was calculated with no initial data transformation and ASVs that were not present in more than 0.1% relative abundance in any sample have been removed. The principal coordinates analysis was based on Bray-Curtis distances. The effect of the route of inoculation on individual members of the bacterial community was computed with MaAsLin2 and a minimum prevalence threshold of 0.1 and a minimum relative abundance threshold 0.01.Benjamini-Hochberg procedure was applied as a correction method for computing the q-values. All raw FASTQ reads for 16S rRNA gene sequences have been deposited under NCBI accession number:

### Real-time qPCR

Real-time qPCR amplification was performed on DNA extracted from the ceca and litter of broiler chicks as described previously^37^ using a CFX96 Touch Real-Time PCR Detection System (Bio-Rad Inc., Hercules, CA). Reaction mixtures (20 μl) for all assays contained 1X SsoAdvanced Universal SYBR Green Supermix (Bio-Rad Inc., Hercules, CA), 600 nM (each) primers (Supplementary Table 5), and 2 μl of cecal/litter DNA. Calibration curves used for converting qPCR cycle threshold values to gene copies per gram were determined using genomic DNA of *E. coli* or *Salmonella* strains harboring the region or gene of interest

### Statistical analyses

Continuous variables did not meet the assumption of a normal distribution; therefore, non-parametric testing for direct comparisons was performed using Wilcoxon rank sum and signed rank tests, and the Kruskal-Wallis rank sum test was used for one-way analysis of variance tests. Furthermore, continuous variables were log-transformed before any statistical tests were performed. Lastly, principal component analysis was performed on litter and ceca samples using the copy number of genes quantified by qPCR as features. The samples were then projected onto the first two principal components and colored by route of SH challenge to visualize each route’s contribution to the data variance. Pearson correlation coefficients were calculated to examine the correlation between *Salmonella ttr* and *E. coli gapA* gene copies present in ceca and litter. Statistical analyses were performed using R (v 3.4.1).

### Data availability

All raw FASTQ reads including short and long reads for sequenced bacterial genomes are publicly available under NCBI accession numbers: PRJNA683658, PRJNA684578 and PRJNA684580. Shotgun and Hi-C reads are publicly available under NCBI accession number: PRJNA688069 and 16S rRNA gene sequences under NCBI accession number: PRJNA669215. The whole genome assembly for *S*. Heidelberg strains SH-2813-ancestor and ic9b harboring p1ST26 have been made available under NCBI biosample number: SAMN12082795 and DDBJ/ENA/GenBank accession numbers JAEMHU000000000, respectively. The whole genome assembly for *E. coli* strain Ec-FL1-1X and Ec-FL1-2X carrying IncI1 are available under GenBank accession numbers CP066836 and JAFCXR000000000, respectively. Variant call format (VCF) files of identified SNPs/indels and the Linux/Unix shell script used have been deposited in Dryad Digital Repository: https://doi.org/10.5061/dryad.4tmpg4f8d.

## Ethics statement

All animal experiments were approved by the University of Georgia Office of Animal Care and Use under Animal Use Protocol: A2017 04-028-A2.

## Acknowledgements

We are grateful to Marlo Sommers, Jasmine Johnson, Carolina Hall, Jeromey Jackson, and Latoya Wiggins for their logistical and technical assistance. This work was supported by USDA Agricultural Research Service (Project Number: 6040-32000-010-00-D), non-assistance cooperative agreement (58-6040-6-030) between USDA Agricultural Research Service and University of Georgia, Research Foundation, and research service agreement (58-6040-8-035) between USDA Agricultural Research Service and Colorado State University. This work was also partially supported by Colorado State University through startup funds to ZA. BZ was supported by the competence center. The competence center (FFoQSI) is funded by the Austrian ministries BMVIT, BMDW, and the Austrian provinces Niederoesterreich, Upper Austria and Vienna within the scope of COMET-Competence Centers for Excellent Technologies. The Austrian Research Promotion Agency FFG handle the program COMET. This study was supported in part by resources and technical expertise from the Georgia Advanced Computing Resource Center, a partnership between the University of Georgia’s Office of the Vice President for Research and Office of the Vice President for Information Technology. Any opinions expressed in this paper are those of the authors and do not necessarily reflect the official positions and policies of the USDA or the National Science Foundation, and any mention of products or trade names does not constitute recommendation for use. The authors declare no competing commercial interests in relation to the submitted work. USDA is an equal opportunity provider and employer.

## Author’s contributions

A.O., Z.A., K.C. and N.A.C. designed the study. A.O., G.Z., D.E.C. and C.R. performed live-broiler chicken studies. A.O., J.L., G.Z., S.H. and D.C. performed bacteriological analyses, antibiotic susceptibility testing, DNA extraction and qPCR. J.L., G.Z. and A.O. performed Illumina whole genome sequencing. A.O. performed cecal shotgun and Hi-C library preparation. T.L. made 16S rRNA gene libraries and sequencing. B.Z. performed 16S rRNA bacterial community analysis and interpretation. K.I., A.H., and A.O. performed phenotype microarray analyses with the help of J.G. and E.L. A.O., Z.A., M.O.P., J.C.T., S.M.L., R.W., J.L., D.C. and L.L performed bioinformatic analyses and data curation. K.H. and A.O performed *Salmonella* Heidelberg fitness testing and analysis. A.O., J.L. and D.C. isolated multidrug resistant *E. coli* donors from cecal contents with help from M.J.R. A.O., B.Z. and M.E.B. performed statistical analyses. A.O., Z.A., N.A.C., B.Z., R.W., M.P., S.M.L., K.I., G.Z., J.L., K.H. and L.L. drafted the manuscript, which was reviewed and edited by all authors. A.O., Z.A., S.A.A., L.C. and M.W. supervised the study.

## Supplementary Methods

### Isolation of *E. coli* from ceca of broiler chicks

To isolate *E. coli*, we retrospectively screened two cecal slurry from intracloacally challenged chicks on CHROMagar™ plates supplemented with gentamicin (8 ppm) and tetracycline (8 ppm). Frozen vials of cecal contents were thawed on ice and vortexed. For each sample, a 10 μl aliquot was struck for isolation, and a 100 μl aliquot was spread plated on CHROMagar™ with relevant antibiotics. Plates were incubated overnight at 37°C. Blue green and blue-cream colonies were counted as presumptive *E. coli* and colonies was restruck for isolation on CHROMagar™ supplemented with gentamicin and tetracycline. After incubation, 2-3 colonies from each sample were selected to represent different forms of blue green/blue-cream colonies and struck in parallel onto sheep blood agar (SBA) and Eosin Methylene Blue (EMB) (Remel Inc., KS, USA) and incubated overnight. Growth on EMB was the characteristic green metallic sheen after incubation. Colonies from SBA were stored in 30% LB glycerol at −80°C and were used for antimicrobial susceptibility testing and whole genome sequencing.

### Long read sequencing quality control

PacBio sequencing of samples was done using using the Sequel II system. MinION sequencing of samples was done using R9.5 chemistry on a 1D flowcell. Lima (https://github.com/PacificBiosciences/pbbioconda) was used to demultiplex barcoded PacBio samples and to convert the split BAM files into fastq format. BBMap reformat.sh (https://github.com/BioInfoTools/BBMap) was used to randomly subsample generated PacBio reads down to 200X coverage. Porechop v0.2.4 (https://github.com/rrwick/Porechop) was used to demultiplex barcoded MinION samples. Raw Illumina reads were trimmed using Trimmomatic v0.39^1^ (command line parameters: PE ILLUMINACLIP:NexteraPE-PE.fa:2:30:10:3 LEADING:3 TRAILING:3 SLIDINGWINDOW:4:20 MINLEN:36).

### Comparative genomics

To determine IncI1-pST26 consensus, first we used BWA^2^ or Bowtie v. 7.2.1^3^ (implemented in Geneious Prime®) to align raw FASTQ files against a complete circular p1ST26 or p2ST26 and used the binary alignment map file generated for consensus determination in Geneious Prime®. To identify identical IncI1-pST26 plasmids, we performed a BLAST search against the NCBI non-redundant database and selected the top 50 plasmids matching p2ST26 from this study. To construct a whole plasmid based maximum likelihood tree, we downloaded the FASTA files for the top 50 plasmids from NCBI and used them for whole genome alignment.

### Tools used for visualizing and exploring high-throughput sequence data

Gene presence/absence heatmap, with gene presence/absence data obtained from Roary, was generated using the pheatmap v1.0.12, tidyverse v1.3.0 and viridis v0.5.1 packages in R v4.0.2. BLAST Ring Image Generator^4^ was used for genome comparison visualization including GC skew change and Phandango^5^ was used for visualizing phylogenetic trees with their associated metadata. PHYLOViZ 2.0^6^ was used to generate a minimum spanning tree of SH isolates from core genome information obtained through SISTR (i.e., wgMLST_330:complete-alleles, wgMLST_330:missing-alleles, wgMLST_330:partial-alleles,closest-public-genome-alleles-matching and cgMLST_cluster_level). SnapGene® was used for drawing linear maps of plasmid, prophages, and other regions of interest in bacterial genomes. Geneious Prime® and SnapGene® were used to view MAFFT alignments of DNA and amino-acid sequences.

---

**Supplementary Table 1:** Assembly report for a *S*. Heidelberg strain with sequencing bias

| Assembly statistics* | Short read<sup>†</sup> | Long read<sup>‡</sup> | Hybrid<sup>§</sup> |
| --- | --- | --- | --- |
| Number of contigs | 554 | 4 | 33 |
| Total length (bp) | 4314964 | 4703397 | 4865484 |
| Reference length (bp) | 4788855 | 4788855 | 4788855 |
| Number of misassembled contigs | 4 | 0 | 1 |
| Misassembled contigs length (bp) | 47430 | 0 | 4742328 |
| Genome fraction of reference (%) | 89.969 | 98.247 | 99.815 |
| Largest alignment (bp) | 60767 | 2669051 | 3233973 |
| Total aligned length (bp) | 4308655 | 4703397 | 4826756 |

\* Assembly was done using Unicycler and SH-ancestor was used as the reference genome (4,788,855 bp)
<sup>†</sup> Illumina short technology was used for sequencing.
<sup>‡</sup> PacBio long read sequencing technology was used.
<sup>§</sup> Illumina short reads and PacBio long reads were combined.

**Supplementary Table 2.** Mutations found in p1ST26 but absent in p2ST26

| Loci* | Location<sup>†</sup> | Product | Mutation<sup>‡</sup> |
| --- | --- | --- | --- |
| 20998 | Intergenic | Upstream replication protein | N/A |
| 23599 | CDS | *yacA* | Substitution |
| 23637 | CDS | *yacA* | Substitution |
| 23661 | CDS | *yacA* | Substitution |
| 23695 | CDS | *yacA* | Substitution |
| 23698 | CDS | *yacA* | Substitution |
| 23704 | CDS | *yacA* | Substitution |
| 23731 | CDS | *yacA* | Substitution |
| 23734 | CDS | *yacA* | Substitution |
| 23826 | CDS | *parE* | Substitution |
| 25883 | CDS | *tniA* | Insertion |
| 35066 | CDS | *aac3-Via* | Substitution |
| 35608 | CDS | *aadA1* | Substitution |
| 35635 | CDS | *aadA1* | Substitution |
| 38685 | CDS | *tn21* | Substitution |
| 41737 | Intergenic | downstream tn3; upstream hypothetical | N/A |
| 42362 | CDS | hypothetical | Substitution |
| 49072 | CDS | hypothetical | Substitution |
| 70253 | CDS | hypothetical | Substitution |
| 70287 | CDS | hypothetical | Substitution |
| 70333 | CDS | hypothetical | Substitution |
| 70343 | CDS | hypothetical | Substitution |
| 92634 | CDS | *icmG/dotF* | Substitution |

\*A circular and complete Inc1 plasmid from Trial 2 was used as the reference genome (111, 898 bp).
<sup>†</sup>CDS- Coding DNA sequence
<sup>‡</sup>N/A – Not applicable

**Supplementary Table 3.**
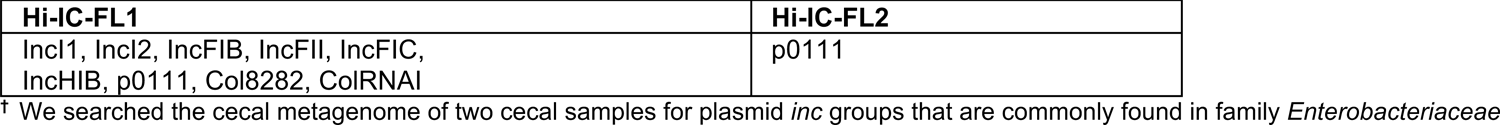
Plasmids found in the cecal metagenome

| Hi-IC-FL1 | Hi-IC-FL2 |
| --- | --- |
| Incl1, Incl2, IncFIB, IncFII, IncFIC,<br>IncHIB, p0111, Col8282, ColRNAI | p0111 |

<sup>†</sup> We searched the cecal metagenome of two cecal samples for plasmid *inc* groups that are commonly found in family *Enterobacteriaceae*

**Supplementary Table 4:** Concentration of metals and pollutants in starter feed.

| Starter feed* | Parts per million (ppm) | Percent (%) | Pounds per ton |
| --- | --- | --- | --- |
| Total Acid Digestion |  |  |  |
| Antimony (Sb) | <0.4 | negligible | negligible |
| Arsenic (As) | 0.999 | negligible | negligible |
| Beryllium (Be) | <0.05 | negligible | negligible |
| Cadmium (Cd) | 0.109 | negligible | negligible |
| Chromium (Cr) | 2.74 | 0.0003 | 0.01 |
| Copper (Cu) | 7.96 | 0.0008 | 0.02 |
| Lead (Pb) | 0.326 | negligible | negligible |
| Nickel (Ni) | 1.67 | 0.0002 | 0 |
| Selenium (Se) | 0.985 | negligible | negligible |
| Silver (Ag) | <0.05 | negligible | negligible |
| Thallium (Tl) | <0.4 | negligible | negligible |
| Zinc (Zn) | 82.7 | 0.0083 | 0.17 |

\*Unmedicated starter feed was synthesized by University of Georgia's poultry research center's feed mill.

**Supplementary Table 5.**
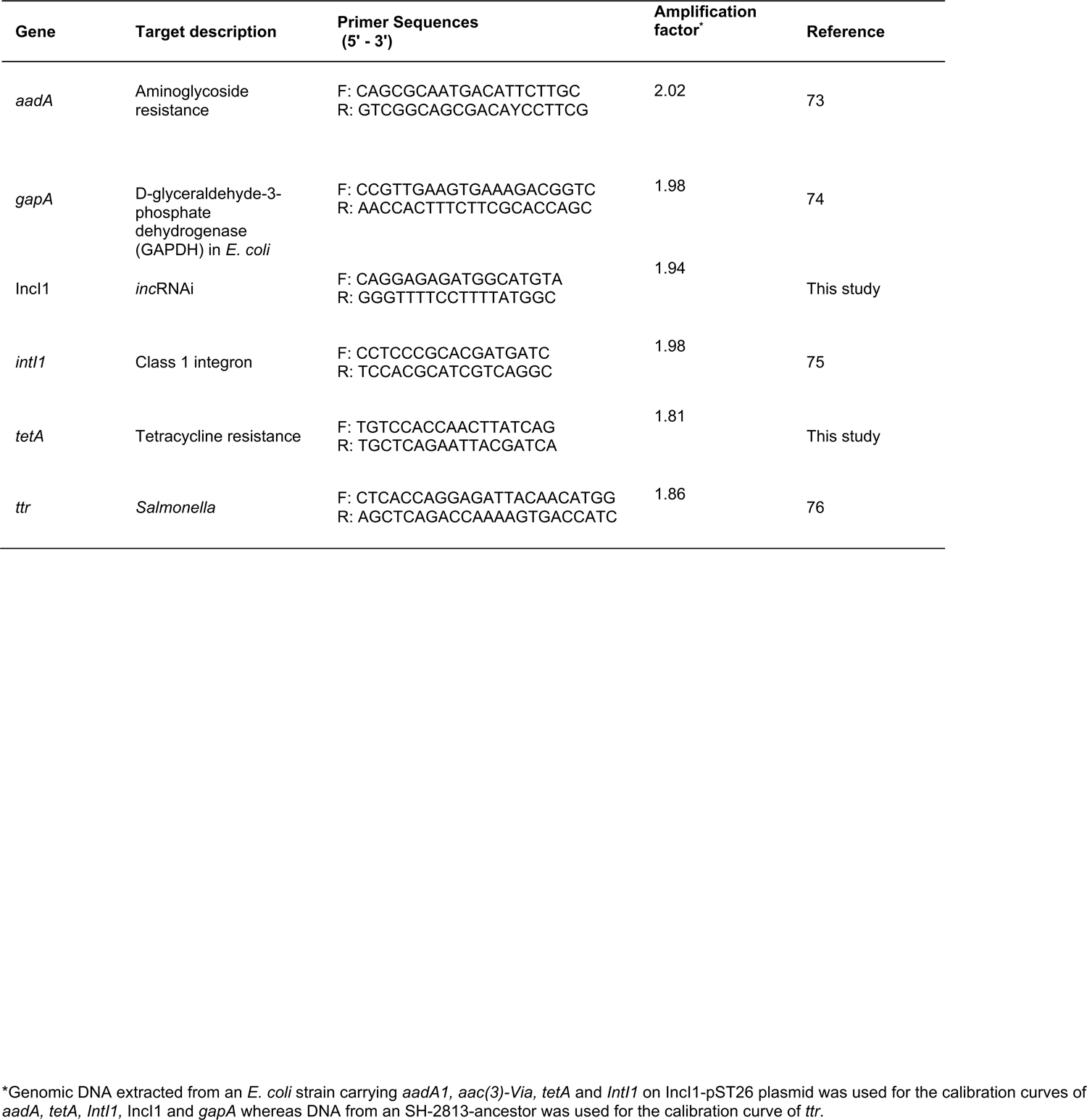
Primers used for the qPCR analysis of ceca and litter.

**Extended Data Fig. 1.**
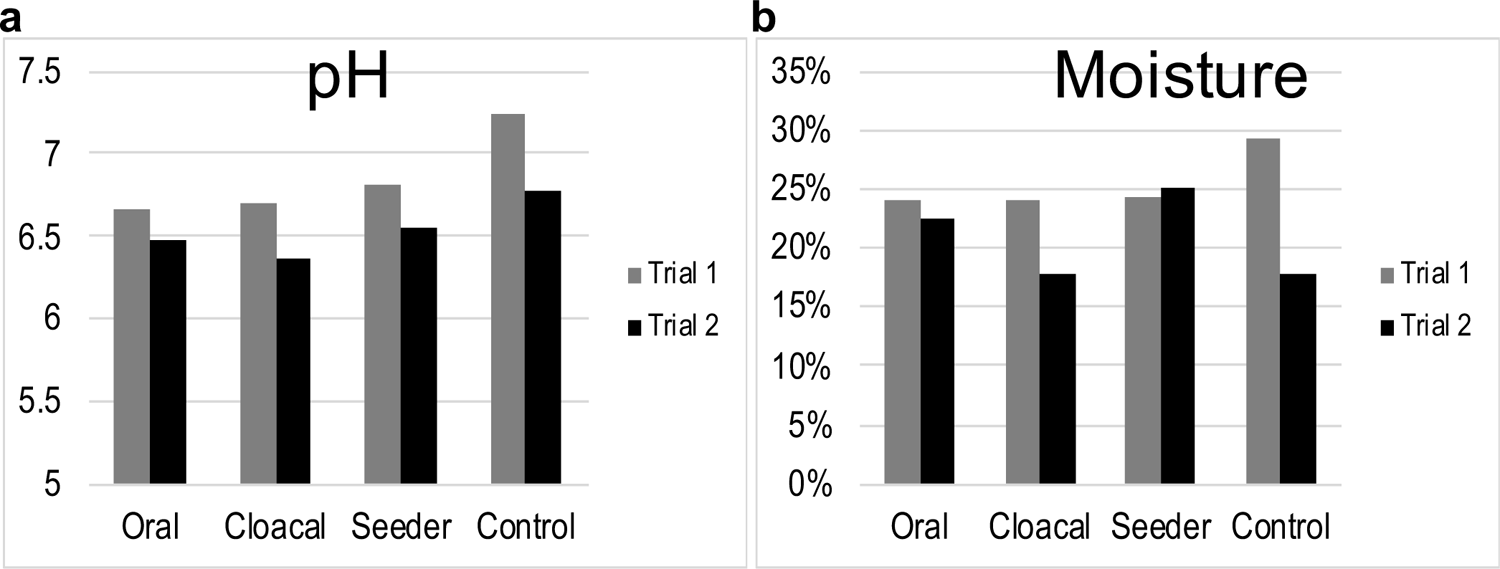
**pH and moisture levels of pen litter**. Litter was collected as grab samples (n = 7) and pooled into one whirl pak bag (n = 1 per pen). The pH of the litter was determined from the eluate made from a 1:5 dilution of litter in 1X PBS.

**Extended Fig. 2.**
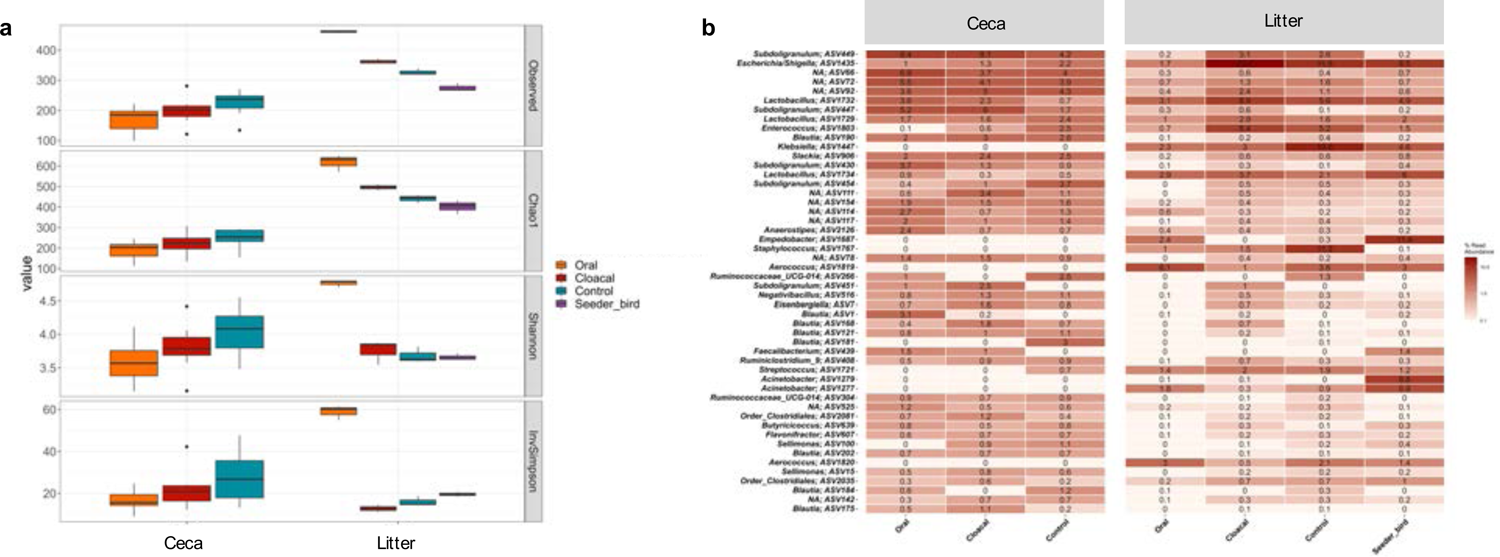
Challenging broiler chicks with SH affected the diversity of bacterial species in their ceca and litter. a, Average alpha diversity indices of ceca and litter samples grouped by route of SH infection. Boxes indicate the interquartile range (75th to 25th) of the data. Whiskers extend to the most extreme value within 1.5 * interquartile range and dots represent outliers beyond that range. d, Heatmap of the top 50 most abundant amplicon sequence variants (ASV) grouped by route of SH infection and split by sample type (Ceca, Litter). Data represents average of ASV counts from replicate 16S rRNA gene libraries for each category.

**Extended Data Fig. 3.**
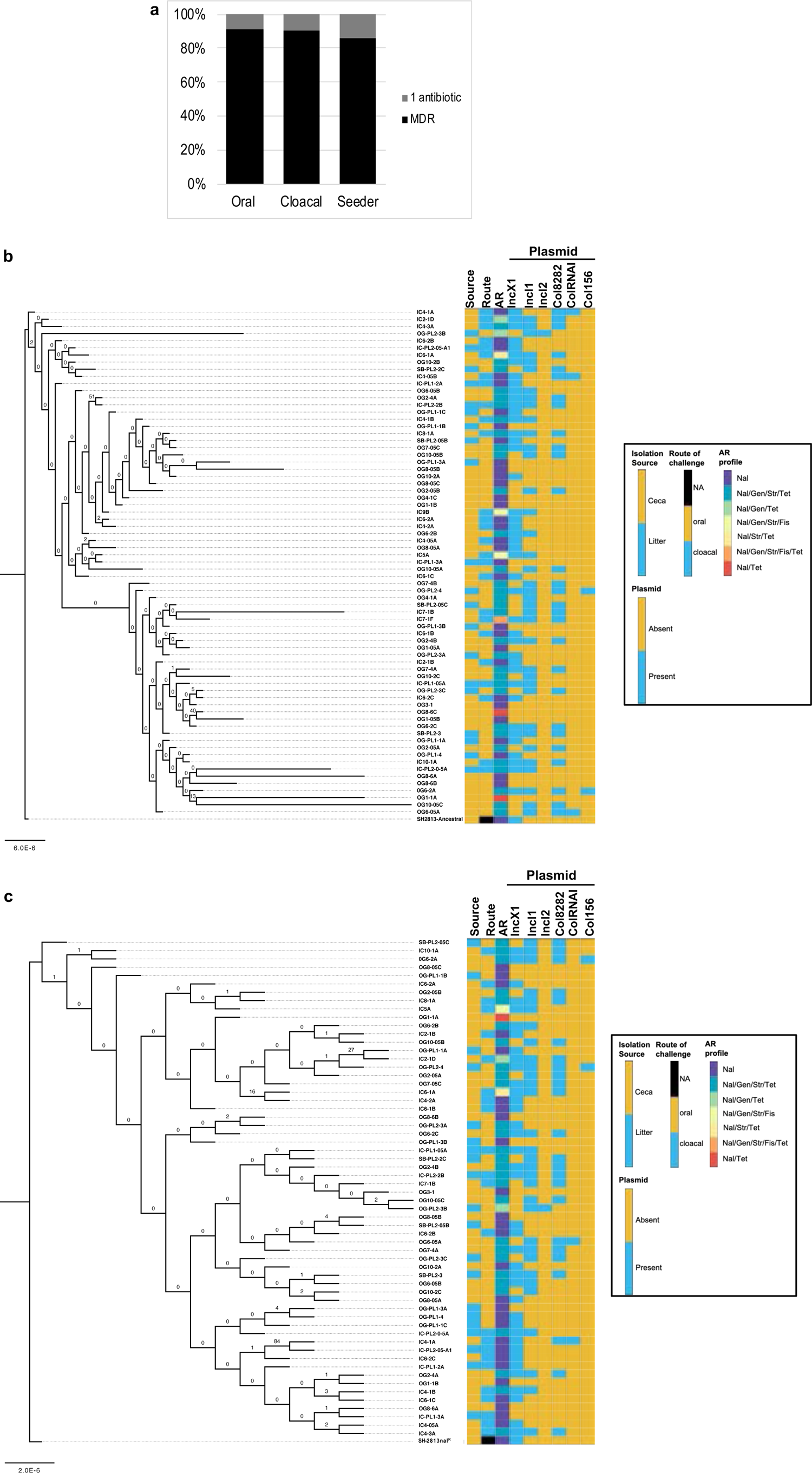
Acquisition of IncI1 plasmid conferred multidrug resistance on SH cells and influenced the contents of their genomes. Antimicrobial susceptibility and whole genome sequencing were performed on SH isolates recovered from the ceca and litter of broiler chicks 2-weeks after challenge. **a**, Percentage of SH isolates (n = 63) from Trial 2 that acquired resistance to two or more antibiotics compared to isolates resistance to only one antibiotic class. Core genome (**b**) and SNP (**c**) based maximum likelihood tree of SH isolates sequenced from Trial 1 (n =2) and Trial 2 (n=69). Isolates with duplicated SNPs/indels were removed before SNP based ML tree was reconstructed. GTR and JC model of nucleotide substitution was used for core and SNP tree, respectively and the GAMMA model of rate heterogeneity were used for sequence evolution prediction. Numbers shown next to the branches represent the percentage of replicate trees where associated isolates cluster together based on ∼100 bootstrap replicates. Tree was rooted with the susceptible SH ancestor for core genome and nal resistant SH ancestor (SH-2813nalR) for SNP tree. (Legend: Nal – nalidixic acid, Gen – gentamicin, Str – streptomycin, Tet – tetracycline, Fis – sulfisoxazole, NA – Not applicable).

**Extended Data Fig. 4.**
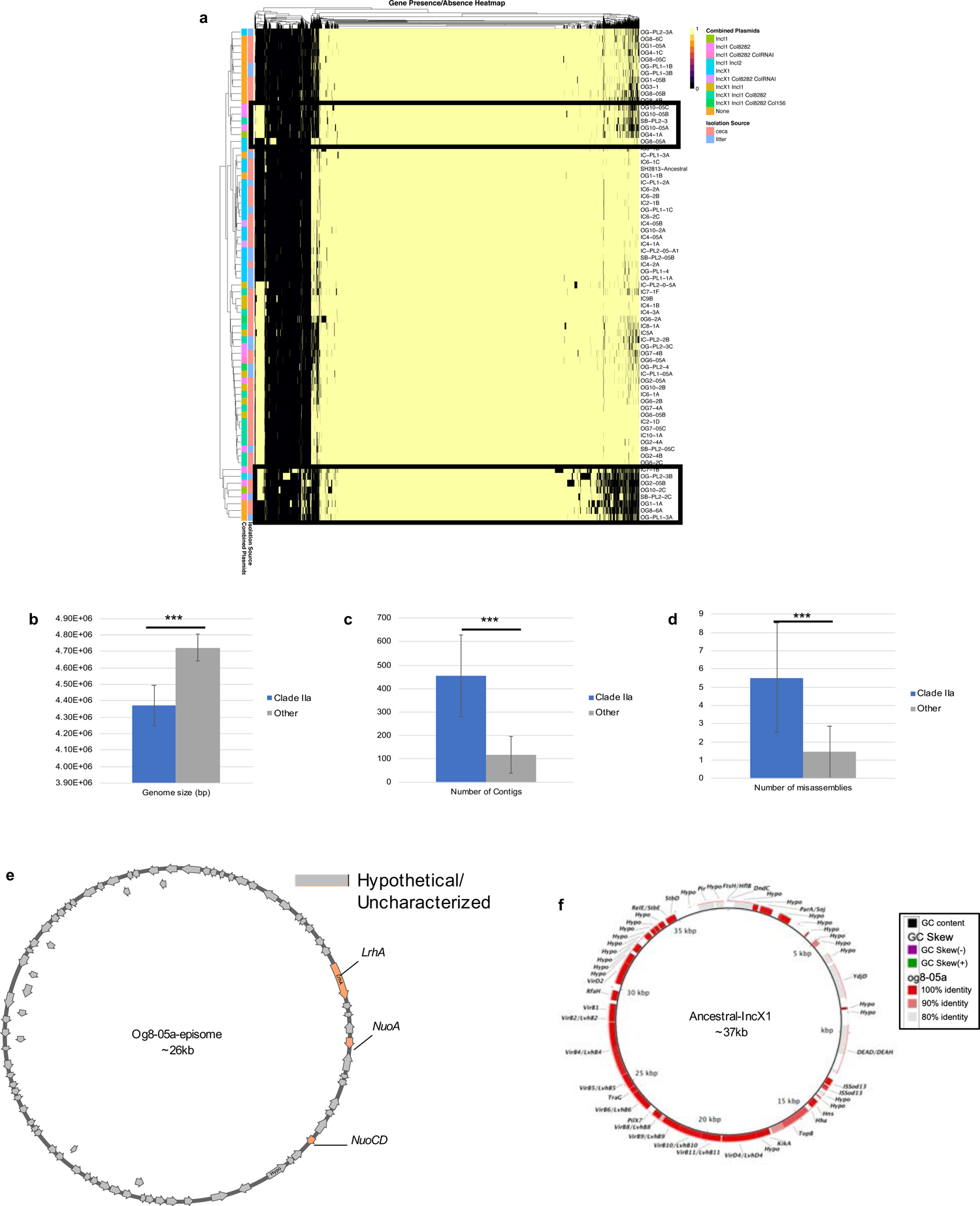
Gene gain and loss contributed to SH diversity in the ceca and litter of broiler chicks a, Heatmap of genes present and absent in SH genomes determined by pangenome analysis. Thick rectangular box highlights SH strains that exhibited higher gene flux and strains with no plasmids but were grouped with SH strains carrying IncI1. **b,c,d** SH genome assembly statistics for SH strains with higher gene flux (n = 12; clade IIa of ML tree reconstructed using accessory genes (see Fig. 2a)) compared to other SH (n=59). (\*\*\**P* < 0.001 (Binomial logistic regression; Error bar = Standard deviation). **e**, Circular map of novel episome present in SH strain og8-05a. **f**, BLASTn alignment of IncX1 present in SH ancestor and IncX1 present in SH strain og8-05a.

**Extended Data Fig. 5.**
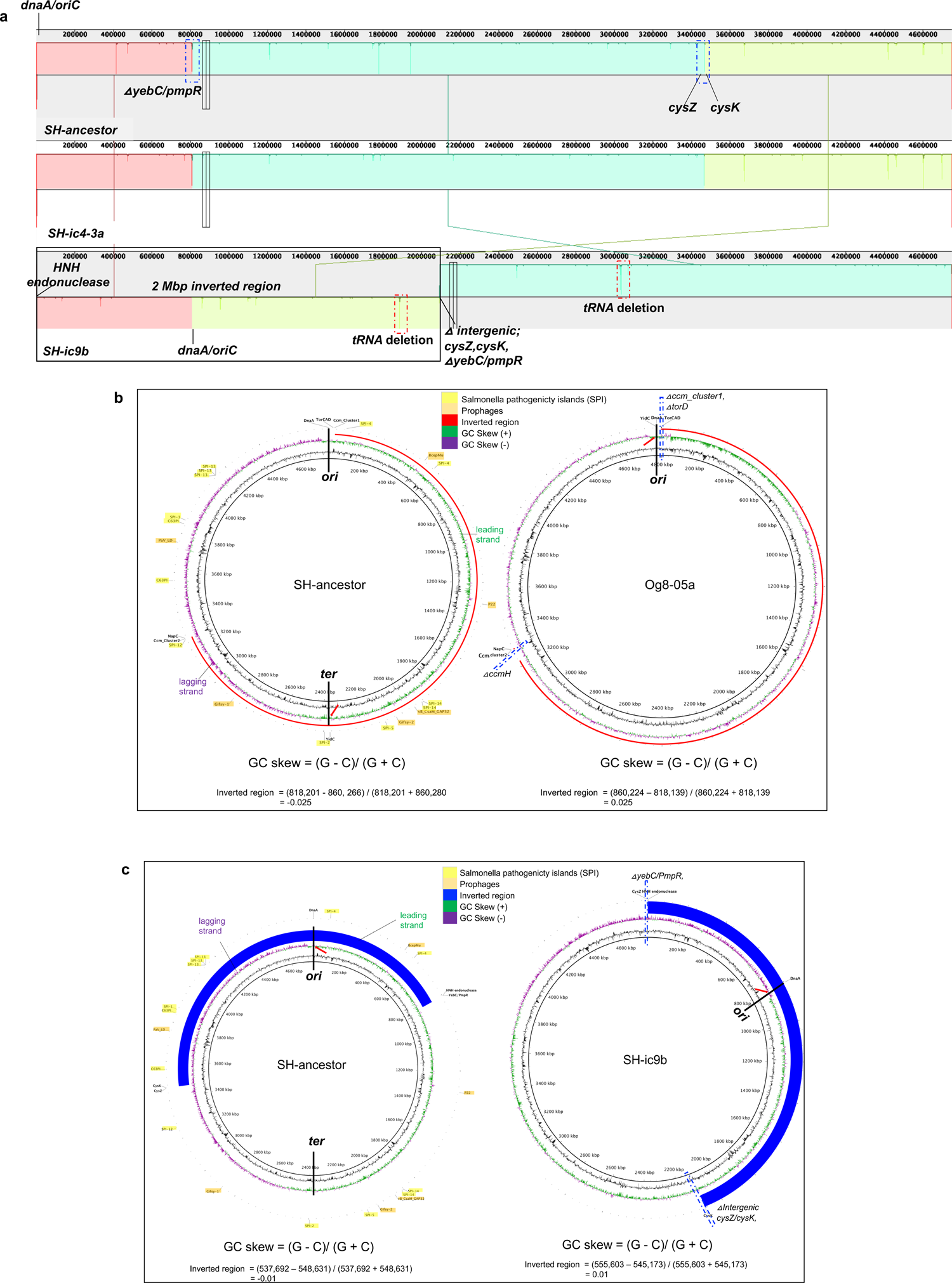
Chromosomal inversion re-oriented the origin of replication of SH strains and positively skewed their GC. Complete chromosome contigs from three SH strains recovered from ceca during Trial 1 (ic9b) and 2 (ic4-3a and og8-05a) were aligned and compared to the ancestor. a, ProgressiveMauve alignment of SH strains ic4-3a and ic9b. A colored similarity plot is shown for each genome, the height of which is proportional to the level of sequence identity in that region. When the similarity plot points downward it indicates an alignment to the reverse strand of the genome of SH-ancestor i.e., inversion. Segment highlighted with solid black rectangular box denotes the inverted 2Mbp region in ic9b. Segment highlighted with horizontal blue box denotes regions with relevant mutations. b and c, GC skew in og8-05a and ic9b compared to SH-ancestor. Red arrow points towards reoriented oriC or ter, whereas red and blue arcs show the inverted region. The number of guanine and cytosine in inverted area was used for GC skew calculation. Dashed blue and red rectangular boxes denotes segments with mutations.

**Extended Data Fig. 6.**
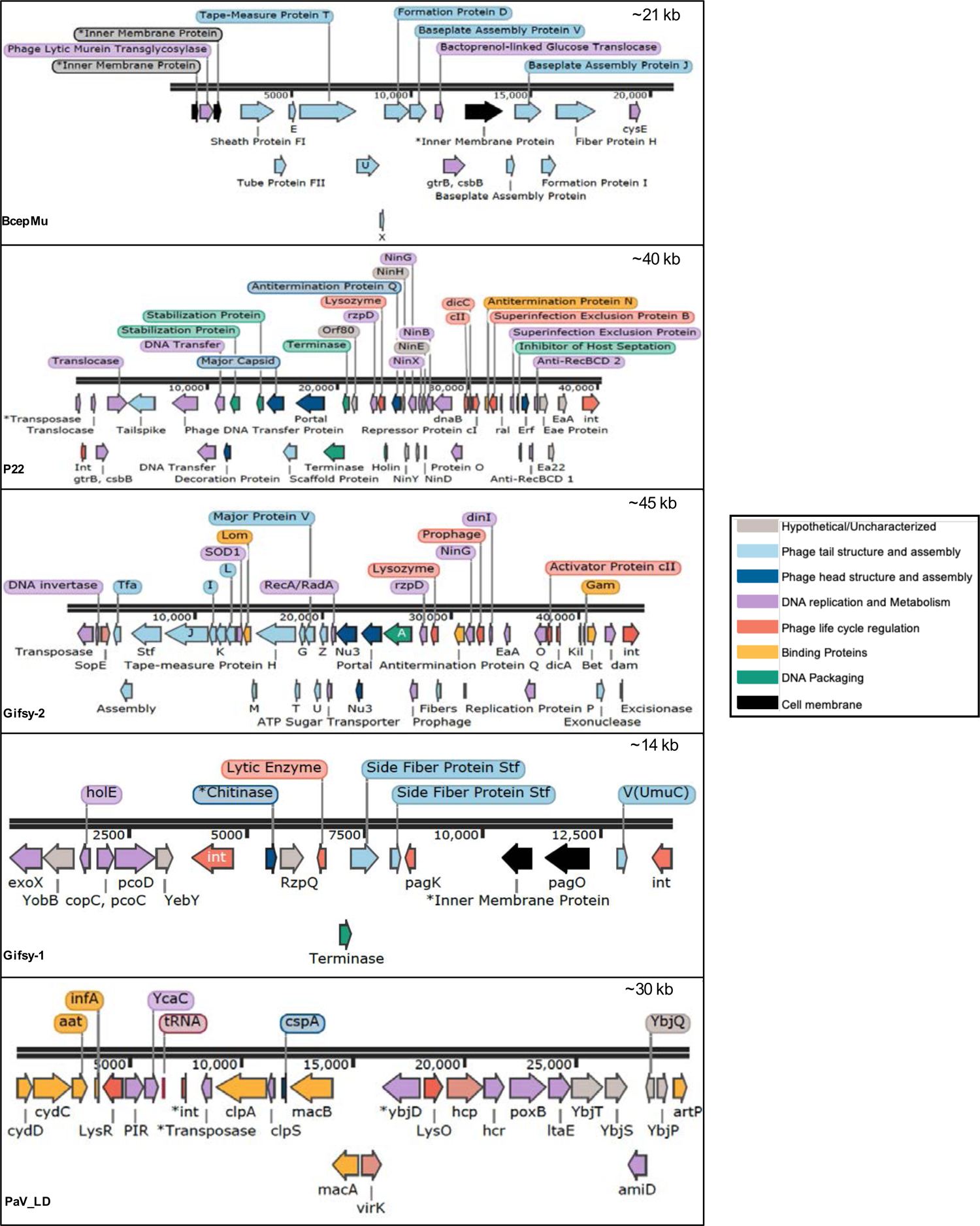
Linear maps of prophages found in SH ancestor.

**Extended Data Fig. 7.**
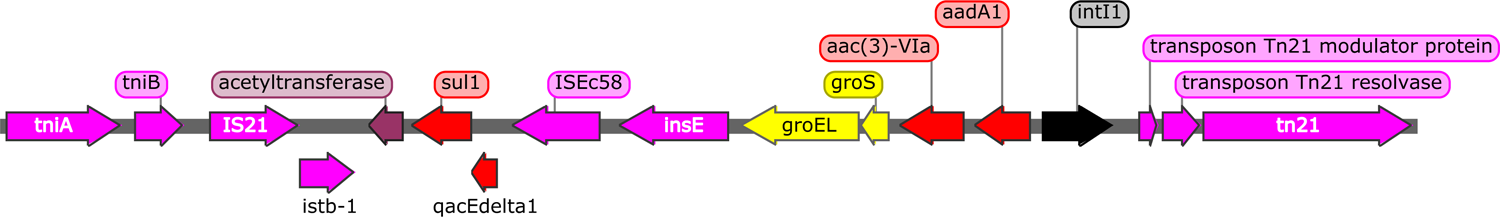
Linear map of the variable region encoding antimicrobial resistance genes in IncI1. The ∼19kb DNA region of the p1ST26 plasmid (∼113kb) present in one multidrug resistant SH isolate from Trial 1 was used for drawing the map.

**Extended Data Fig. 8.**
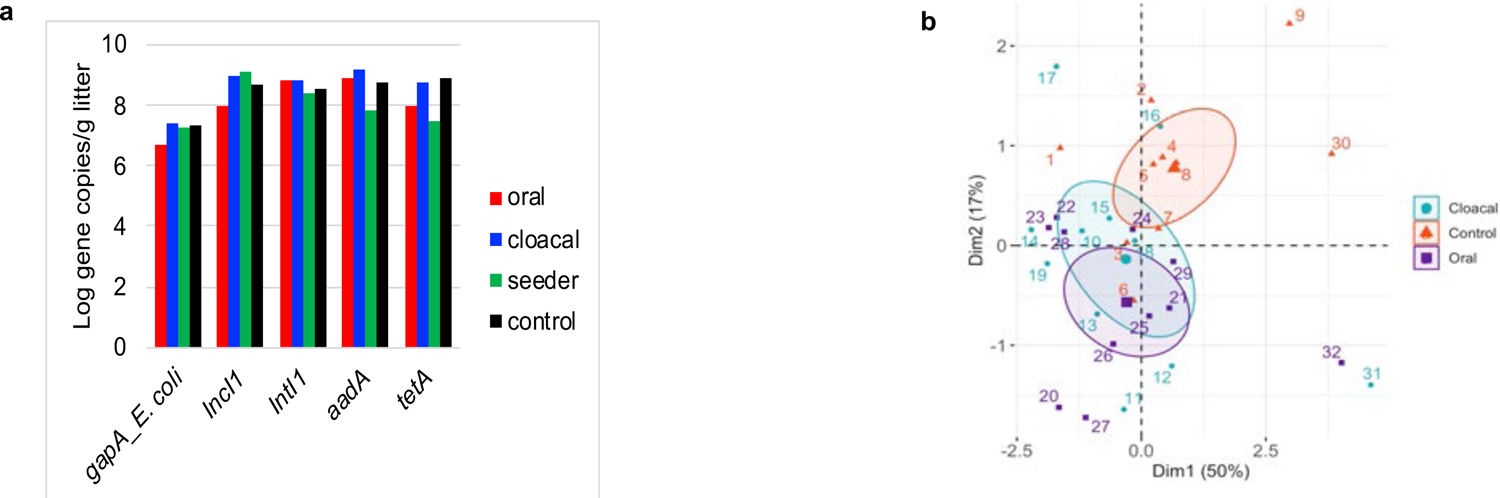
SH challenge affected the abundance of antimicrobial genes a, Concentration of *E. coli* and resistome (IncI1, IntI1 and antibiotic resistance genes) present in the litter of broiler chicks. Gene copies was determined using qPCR. b, Principal component analysis (PCA) characterizing the differences between samples grouped by route broiler chicks got challenged with SH. The PCA was constructed using the concentration of *E. coli, Salmonella* and resistome genes present in ceca and litter of broiler chicks challenged with SH and determined using qPCR. Each number and dot denote one sample, and data is from 32 samples. The farther apart two samples are from each other, the more different they are. Ellipses represents the mean/confidence interval if all the dots of the same color are represented by just one sample.

**Extended Data Fig. 9.**
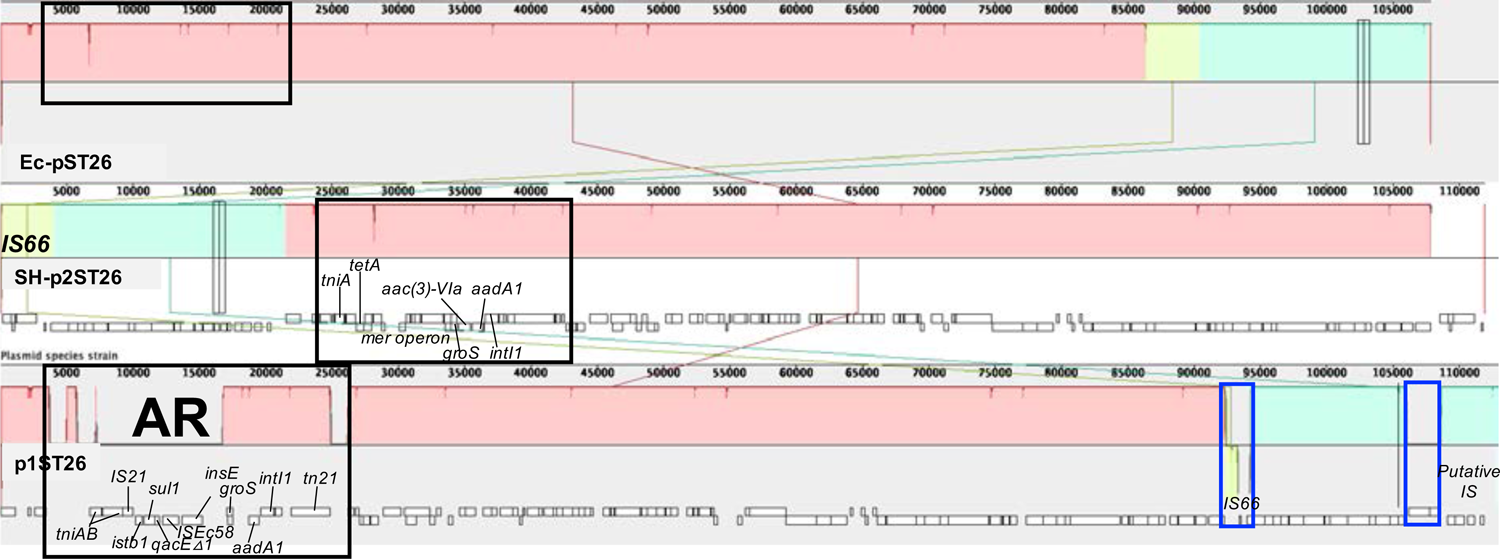
**IncI1-pST26 acquired by SH are identical to the pST26 present in commensal *E. coli* strains**. Complete pST26 contigs from two SH and one *E. coli* strain recovered from the ceca of broiler chicks were used for ProgressiveMauve alignment. A colored similarity plot is shown for each genome, the height of which is proportional to the level of sequence identity in that region. When the similarity plot points downward it indicates an alignment to the reverse strand of the genome of Ec-pST26 i.e., inversion. Segment highlighted with solid black rectangular box denotes the ∼ 20kb region encoding antimicrobial resistance (AR), whereas blue horizontal box denotes region encoding transposons.

**Extended Data Fig. 10.**
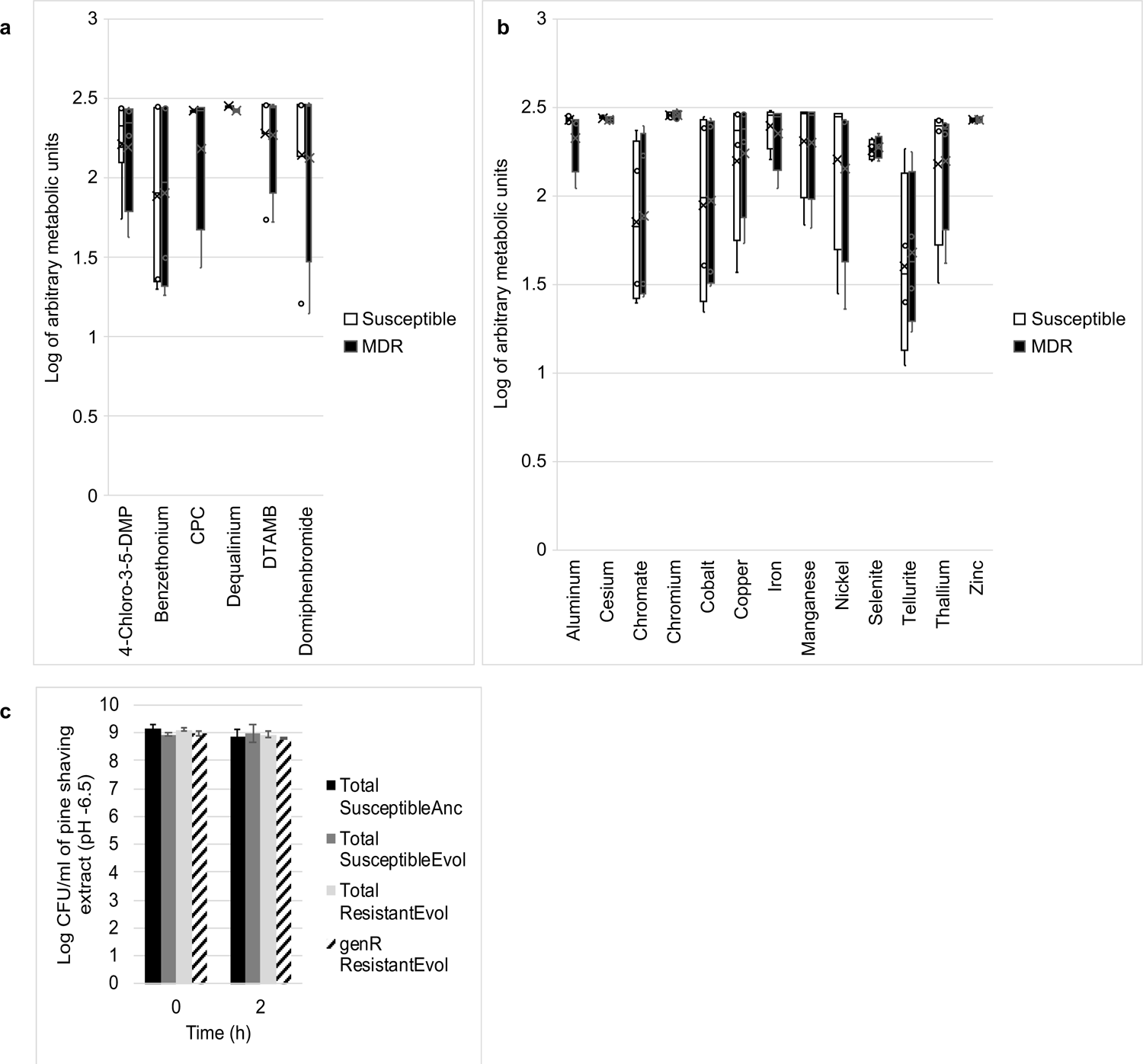
**Exposing SH to metal chlorides, disinfecting compounds or neutral pH did not significantly change their metabolism**. **a,b** Box plots comparing the metabolic activity of one evolved susceptible (no IncI1) and one multidrug (MDR carrying pST26) SH strain using phenotype microarray (PM) plates (Biolog Inc). Pre-configured PM plates are composed of microtiter plates with one negative control well and 95 wells pre-filled with or without nitrogen or amino acid nutrient source and at pH of 3.5 – 10 (n = 1 plate per strain; no of wells per plate per compound was 4) **c,** Concentration of evolved susceptible and MDR SH strains when exposed to pine shaving extract of pH 6.5. Each bar represents the average concentration (colony forming units per ml of pine shaving extract) of three individual population that was established from three single bacterial colonies from three different evolved strains. SH-Ancestor population was established from three single bacterial colonies of SH-2813nal^R^. (Anc = Ancestor, Evol – Evolved i.e., SH isolate recovered during in vivo experiment, genR – population resistant to gentamicin). (Error bars = Standard deviation; *P* > 0.05 (Wilcoxon signed rank test and Kruskal-Wallis rank sum test).

**Extended Data Fig. 11.**
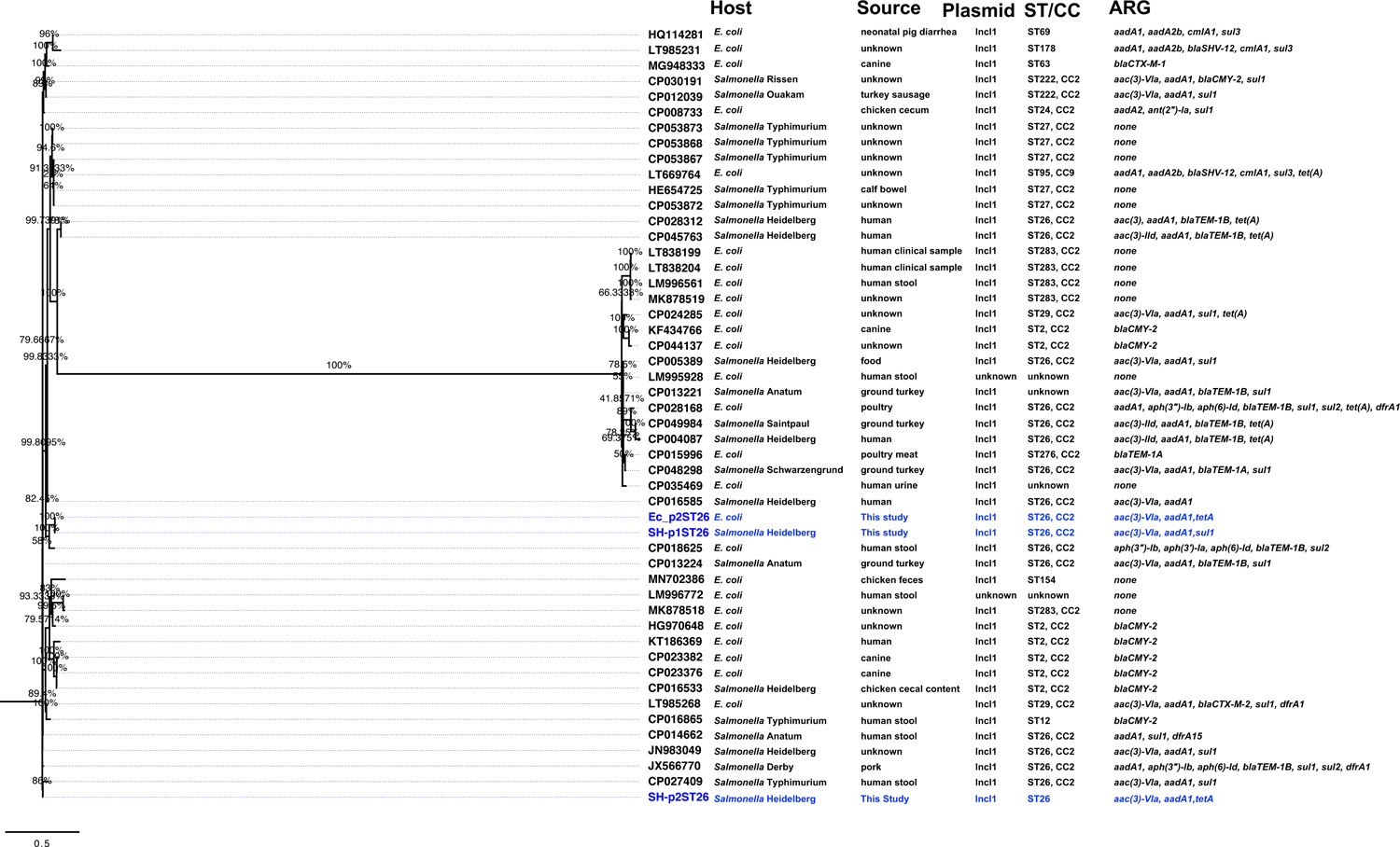
IncI1 plasmids of multilocus sequence type 26 (pST26) are carried by strains of *Salmonella enterica* and *E. coli* isolated from animal and human sources. Maximum likelihood tree reconstructed using the complete DNA sequence of the fifty closest plasmids to pST26 found using NCBI BLASTn search. After multiple alignment was performed, duplicated plasmids (n =4) before ML tree were built. JC model of nucleotide substitution and GAMMA model of rate heterogeneity was used for sequence evolution prediction. Numbers shown next to the branches represent the percentage of replicate trees where associated isolates cluster together based on ∼100 bootstrap replicates. Tree was rooted with p2ST26 plasmid from this study. Taxa in blue text are pST26 plasmids from this study. (Note. ST/CC-Multi locus sequence/clonal complex; ARG – Antibiotic resistance gene)

**Extended Data Fig. 12.**
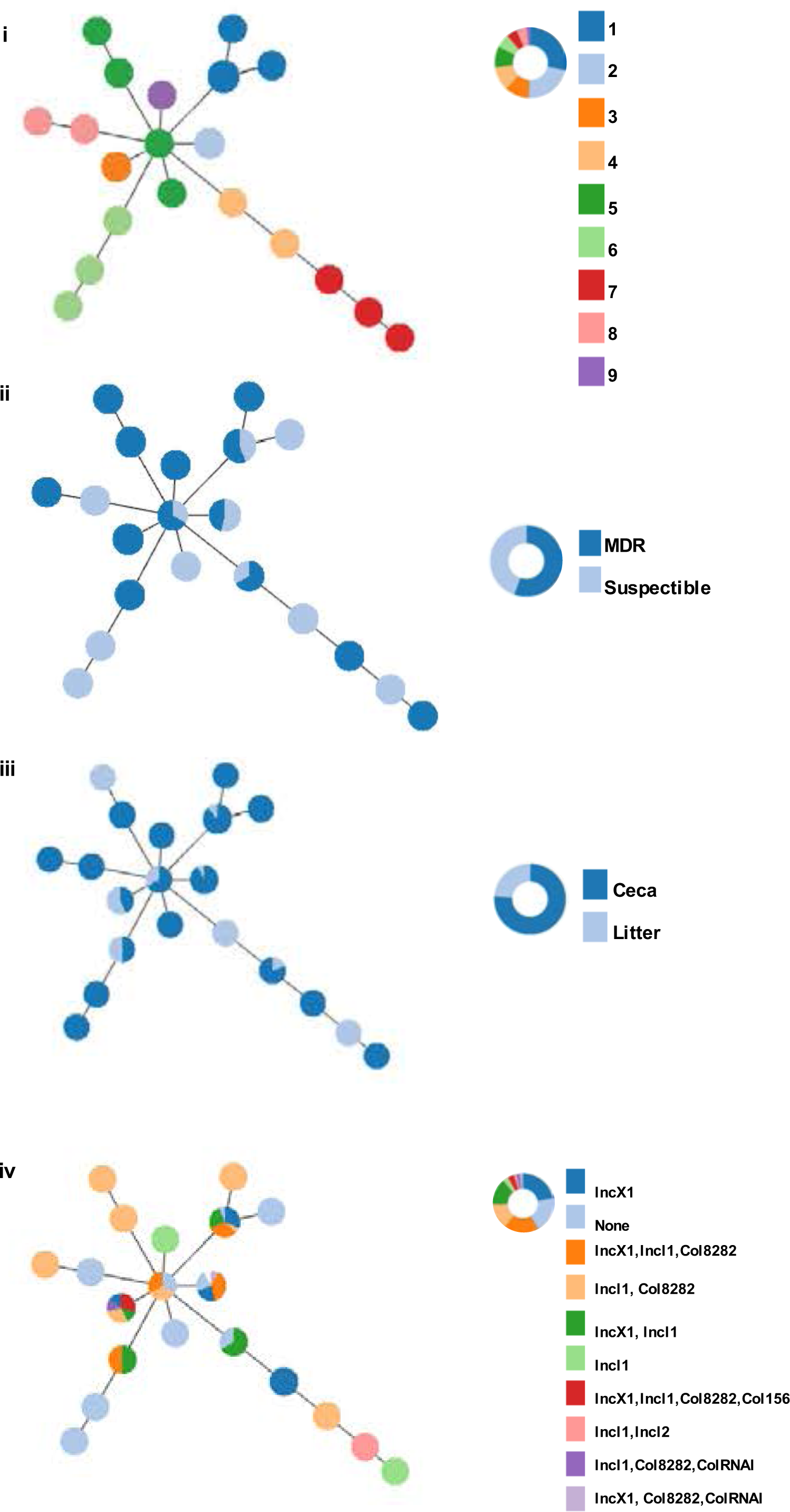
**Minimum spanning tree (MST) of SH genomes from this study**. The core genome multilocus sequence type (cgMLST) data generated by SISTR for 63 genomes from this study including SH-ancestor were used for MST construction. Colors were attributed to nodes according to the number cgMLST cluster levels found (I), antibiotic resistance profile (II), isolation source (III) and plasmid carried. (i) The susceptible SH-ancestor is part of cgMLST cluster 1 (n = 18 isolates). All nodes with distances equal or above 0 were linked. All links from the MST with a distance value above 4 were deleted.

## Supplementary data

**Supplementary data 1:**
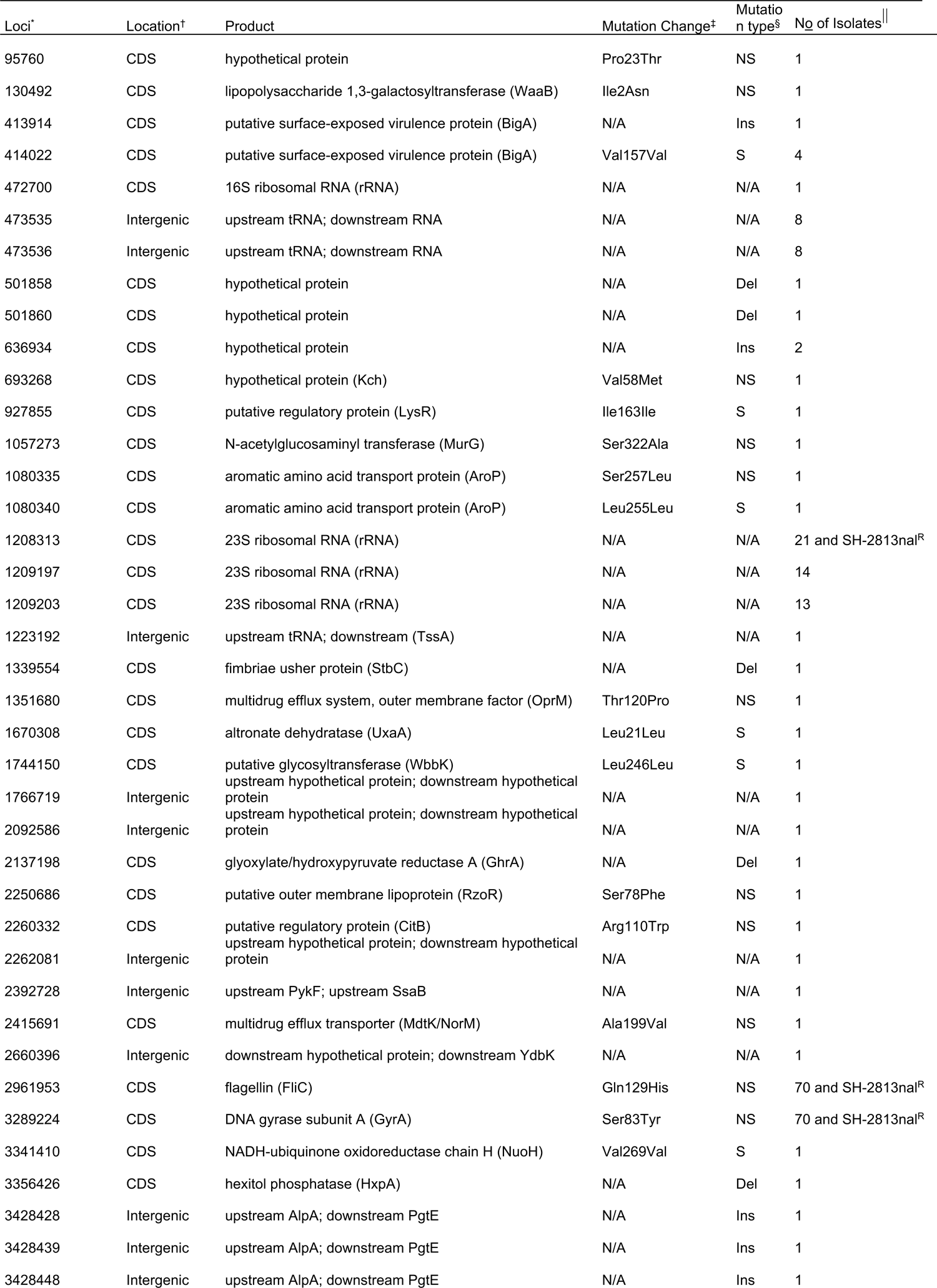
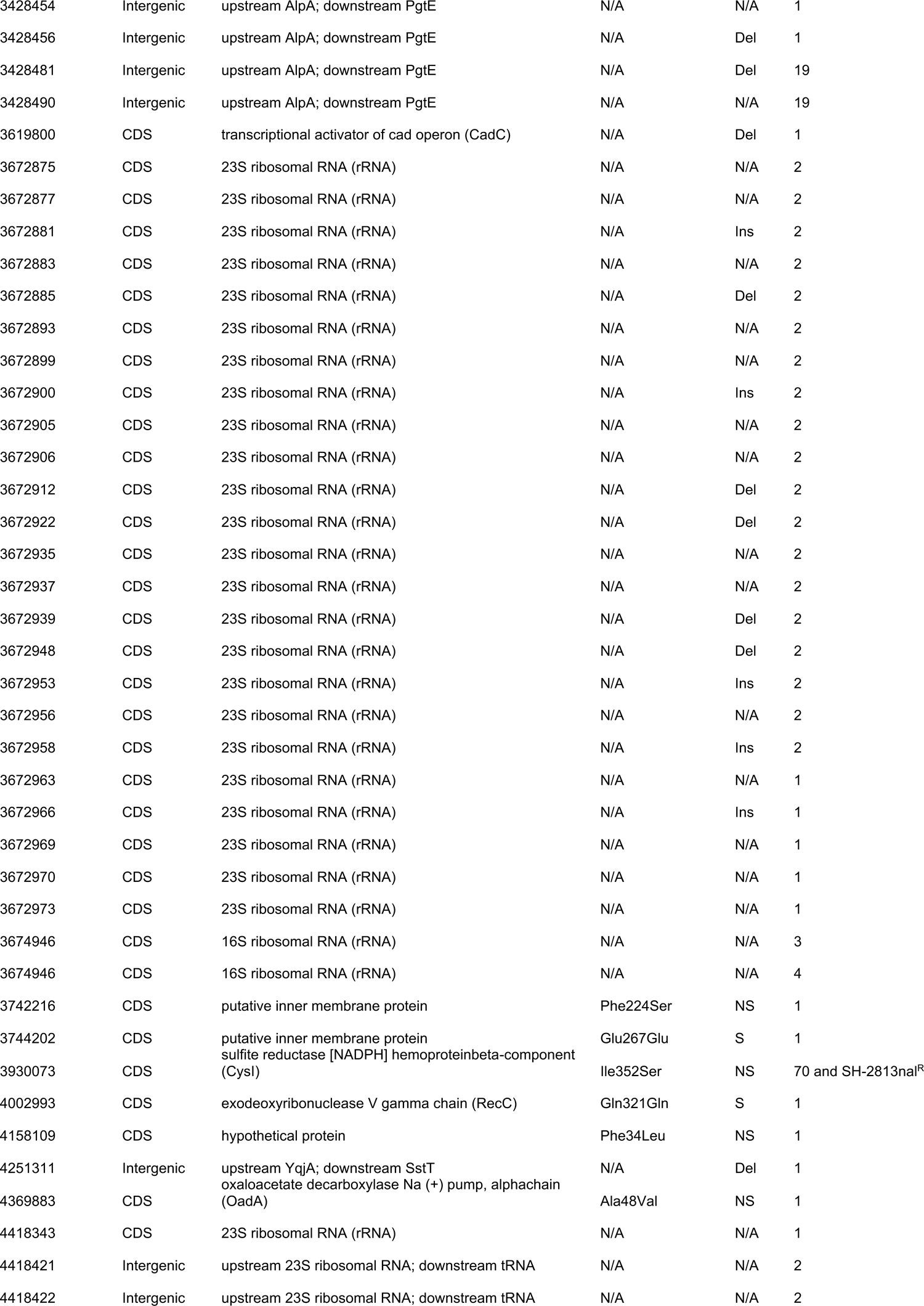
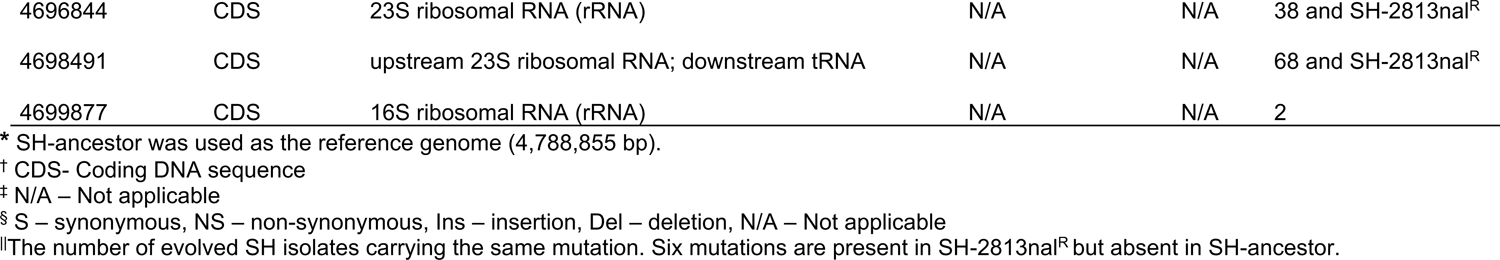
Mutations found in *S*. Heidelberg isolates recovered from ceca and litter.

**Supplementary data 2:**
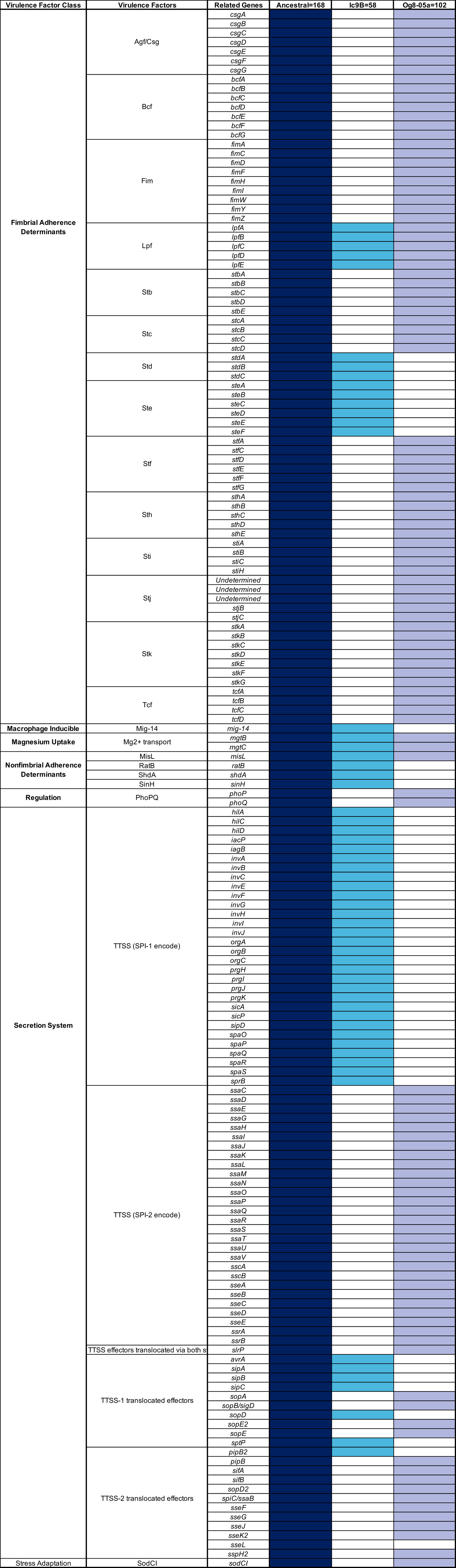
Virulence genes present in SH-ancestor and the inverted region of the genomes of SH strains ic9b and og8-05a. The number of virulence genes found in each genome is written in the column headers.

**Supplementary data 3:**
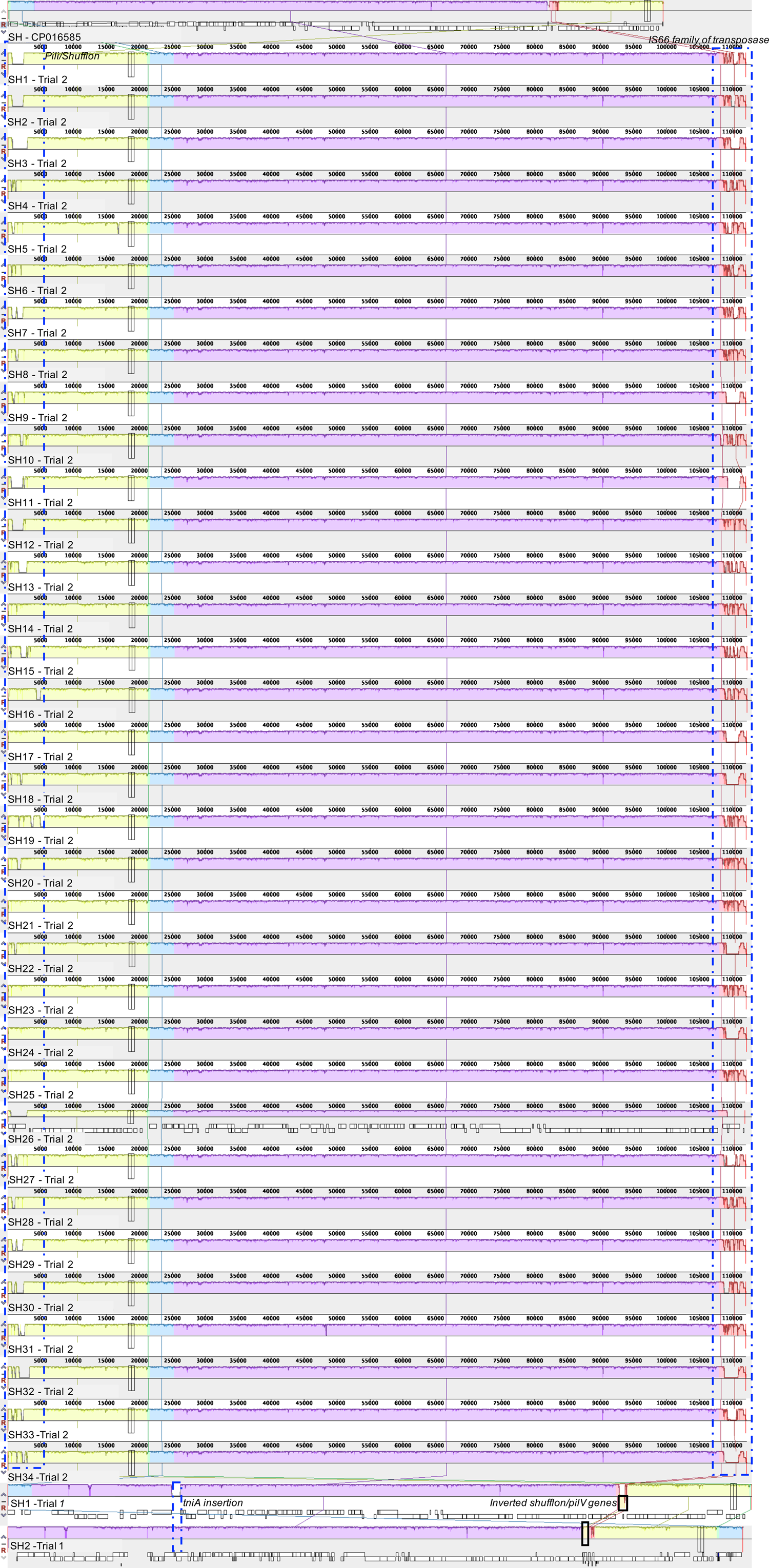
Alignment of IncI1-pST26 plasmids from this study using ProgressiveMauve. Complete plasmid DNA consensus sequence (n= 36) was determined by aligning whole genome FASTQ reads to a complete circular pST26 plasmids from this study. IncI1-pST26 (GenBank: CP016585) present in a *S*. Heidelberg strain available on NCBI was used as a reference genome for the alignment. Segments highlighted with dashed horizontal blue box denotes regions with relevant mutations.

**Supplementary data 4:**
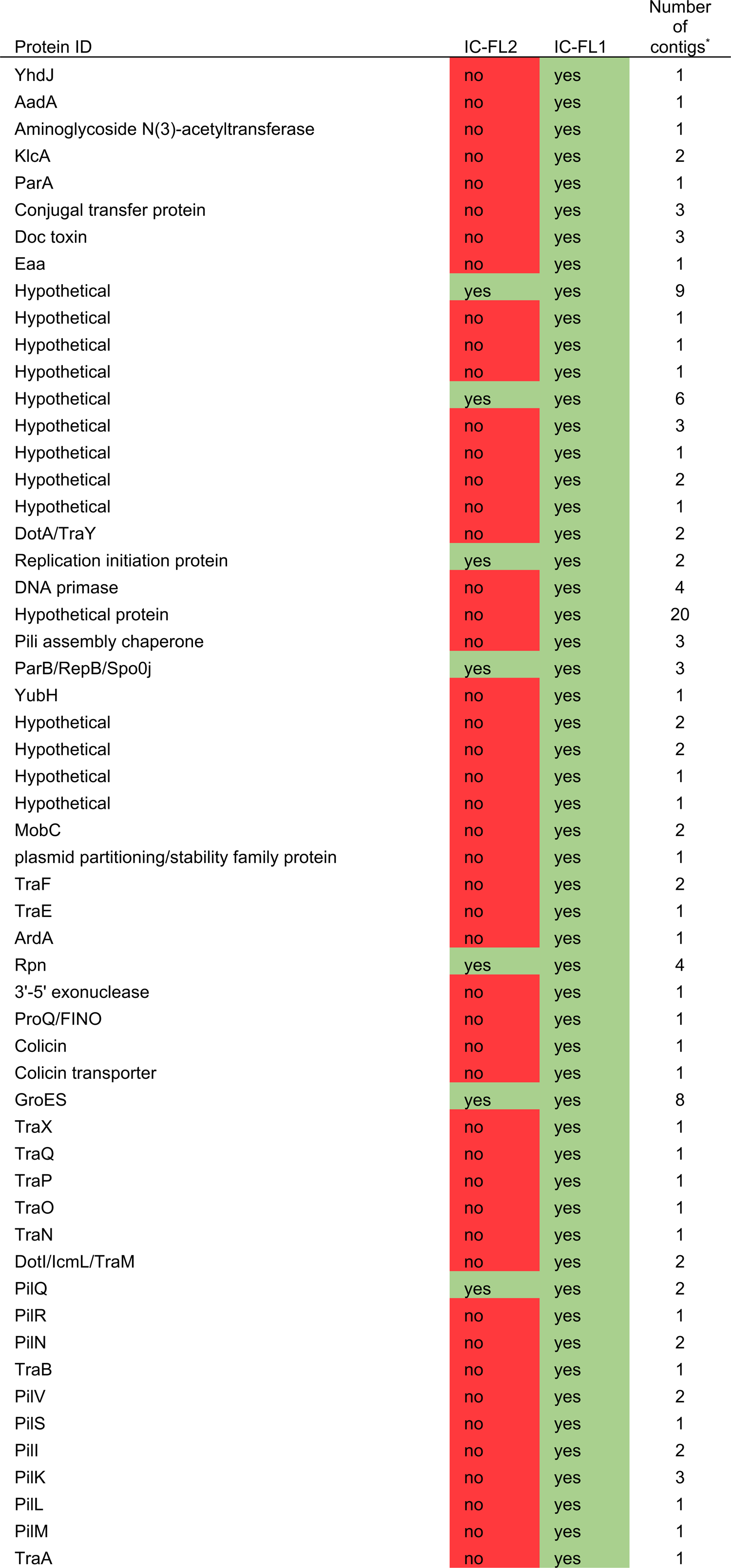
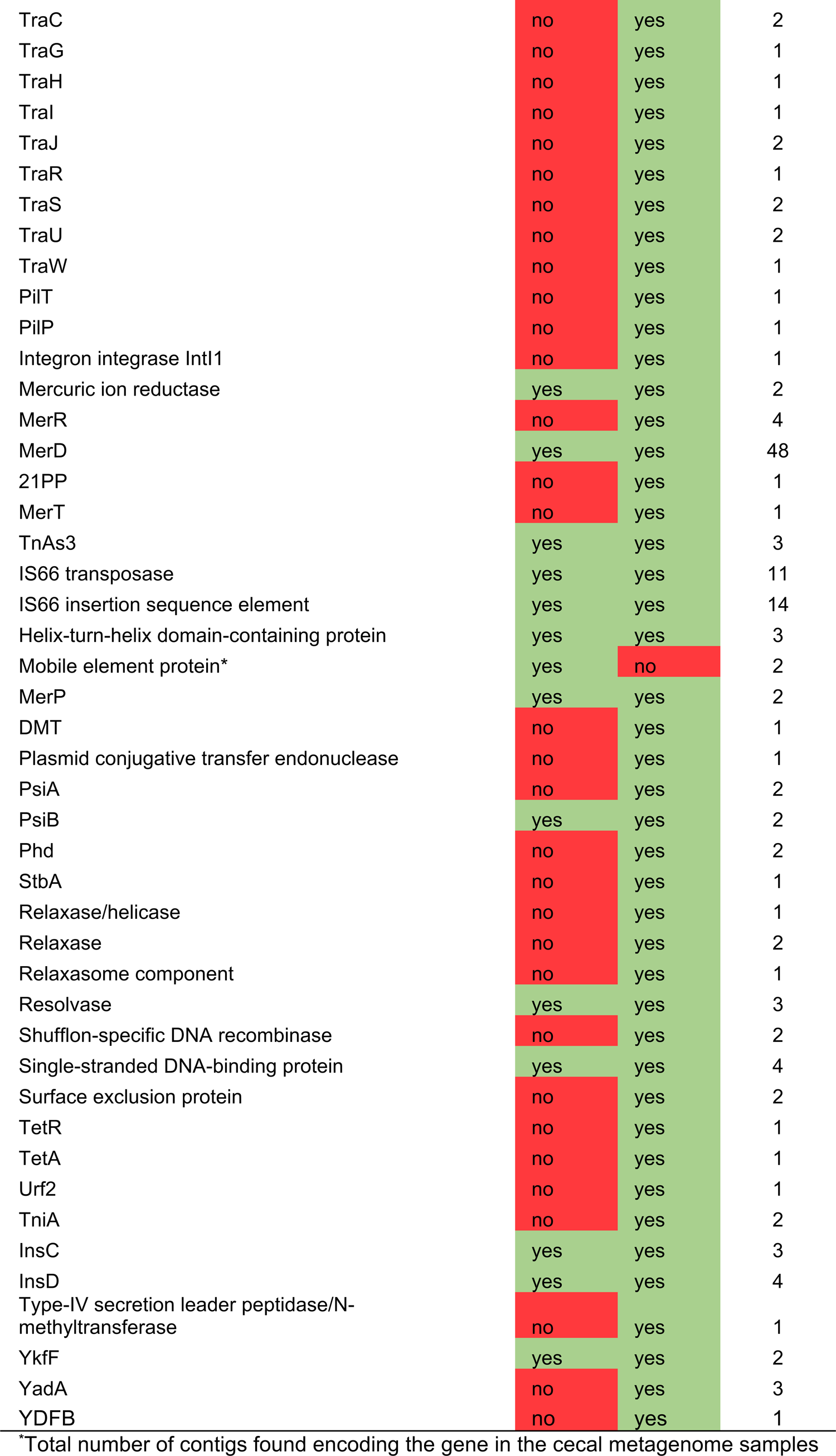
Contigs matching p2ST26 found in the cecal metagenome.

**Supplementary data 5:** Assembly report for bacterial genomes sequenced using short and long reads technology.

